# Ever-increasing viral diversity associated with the red imported fire ant *Solenopsis invicta* (Formicidae: Hymenoptera)

**DOI:** 10.1101/2020.08.11.245274

**Authors:** César A.D. Xavier, Margaret L. Allen, Anna E. Whitfield

## Abstract

**Background:** Advances in sequencing and analysis tools have facilitated discovery of many new viruses from invertebrates, including ants. *Solenopsis invicta* is an invasive ant that has quickly spread around world causing significant ecological and economic impacts. Its virome has begun to be characterized pertaining to potential use of viruses as natural enemies. Although the *S. invicta* virome is best characterized among ants, most studies have been performed in its native range, with little information from invaded areas.

**Methods:** Using a metatranscriptome approach, we further characterized viruses associated with *S. invicta*, in two introduced areas, U.S and Taiwan. The data set used here was obtained from different stages (larvae, pupa, and adults) of *S. invicta* life cycle. Publicly available RNA sequences from GenBank’s Sequence Read Archive were downloaded and *de novo* assembled using CLC Genomics Workbench 20.0.1. Contigs were compared against the non-redundant protein sequences and those showing similarity to viral sequences were further analyzed.

**Results:** We characterized five putative new viruses associated with *S. invicta* transcriptomes. Sequence comparisons revealed extensive divergence across ORFs and genomic regions with most of them sharing less than 40% amino acid identity with those closest homologous sequences previously characterized. The first negative-sense single-stranded RNA viruses included in the orders *Bunyavirales* and *Mononegavirales* are reported. In addition, two positive single-strand viruses and one single strand DNA virus were also characterized. While the presence of a putative tenuivirus associated with *S. invicta* was previously suggested to be a contamination, here we characterized and present strong evidence that Solenopsis invicta virus 14 (SINV-14) is a tenui-like virus that has a long-term association with the ant. Furthermore, based on virus abundance compared to housekeeping genes, phylogenetic relationships, and completeness of viral coding sequences, our results suggest that four of five viruses reported, those being SINV-14, SINV-15, SINV-16 and SINV-17, replicate in the ant *S. invicta*.

**Conclusions:** The present study expands our knowledge about viral diversity associated with *S. invicta* in introduced areas with potential to be used as biological control agents, which will require further biological characterization.

## Background

Insects are the most abundant and diverse group of animals on earth (1). High throughput sequencing has led to huge advances in revealing previously unknown diversity of insect viruses, significantly contributing to filling deep phylogenetic gaps along evolutionary history within the most diverse viral lineages (2–4). Nonetheless, like insect diversity, the diversity of viruses associated with insects is far from clear (1, 3). While many studies have focused on viromes of arbovirus-transmitting insects, especially those involved in transmission of medically important viruses, such as mosquitoes (5–8), other groups that are important pests either impacting agricultural or natural ecosystems, such as invasive ants, have been barely studied (9–11). In addition to contributing to better understanding of basic aspects of virus ecology and evolution, these studies may contribute to new opportunities to use viruses as tools to develop more sustainable insect control methods (12–14).

The red imported fire ant, *Solenopsis invicta*, is an invasive pest causing significant ecological impact and economic loss in invaded areas (15, 16). Originating from South America, *S. invicta* was accidentally introduced into the southern region of the United States (U.S.) almost a century ago, becoming a serious problem (17). Since then, it has spread throughout the southeastern U.S. and more recently into Oklahoma, New Mexico, Arizona, and California (18). Limited introduction events, likely associated with small founder populations, led to a significant reduction in natural enemies and enemy diversity associated with *S. invicta* in introduced areas (19–21). Therefore, *S. invicta* populations may reach sizes even greater than those observed in its native range, making control difficult and even more costly (22). High densities observed in *S. invicta* populations in the U.S. have facilitated its dispersal across the world, contributing to repeated introduction events in several countries, such as China, Taiwan, and Australia (18). Morrison, Porter (23) demonstrated, based on predictive models, that most tropical and subtropical regions worldwide are potentially appropriate for *S. invicta* infestation. Highly competitive ability, generalist feeding habits and high populations densities make *S. invicta* a successful invasive species causing huge disturbance in biodiversity by displacing native ants and other arthropods in introduced regions (15). Currently, chemical insecticides are the most common control strategy used against *S. invicta* (24). Low efficacy due to temporary effects of chemical applications, high cost in extensive areas, and off-target effects harmful to beneficial and other native species are substantial impairments to addressing invasive ant damage and expansion. Therefore, establishment of management strategies that are both environmentally friendly and self-sustainable are necessary.

Classical biological control has been one strategy used in an attempt to control this pest in the U.S., with viruses considered a promising resource to be used as biopesticides (24–26). Over the last decade, a great effort has been made in characterizing the *S. invicta* virome pertaining to potential use in biological control (11, 27–30). To date, the *S. invicta* virome is composed of mainly positive-sense single-strand RNA (+ssRNA) viruses in the order *Picornavirales*. These include eleven viruses from families *Dicistroviridae*, *Polycipiviridae*, *Iflaviridae*, *Soliniviridae* and two unclassified viruses (11, 31). Additionally, one double-strand RNA (dsRNA) and one single-strand DNA (ssDNA) of the families *Totiviridae* and *Parvoviridae*, respectively, have been characterized (11, 32). While most of these viruses have been reported associated with *S. invicta* in its native range, only the species *Solenopsis invicta virus* 1 (SINV-1), SINV-2, SINV-3, SINV-6 and *Nylanderia fulva virus 1* (NfV-1) have been reported in United States (11, 31). Although great progress has been reached in identification and molecular characterization of *S. invicta* virome, the effect of these viruses and their potential use in biological control still need to be elucidated. Interactions among *S. invicta* and SINV-1, SINV-2 and SINV-3, have been the only ones previously characterized (25, 33, 34). SINV-1 was shown to affect claustral queen weight making them lighter than uninfected ones, whereas SINV-2 directly affected fitness of queens by reducing their reproductive output (33). SINV-3 was shown to be the most aggressive virus, causing significant mortality in *S. invicta* colonies, with greatest potential as a biological control agent (25, 35).

Here using a metatranscriptome approach, we further characterized the viruses associated with *S. invicta* in two introduced areas, U.S and Taiwan. These investigations utilized existing publicly available RNA sequences deposited in NCBI GenBank as Sequence Read Archive (SRA) data files (Table 1). Five new viruses were found associated with *S. invicta*, with the first negative-sense single-stranded RNA (-ssRNA) viruses included in the orders *Bunyavirales* and *Mononegavirales* reported. In addition, two +ssRNA viruses, included in the family *Iflaviridae* and an unassigned species, and a partial ssDNA virus, were also characterized.

**Table 1.** Information summary of *S. invicta* transcriptome dataset analyzed in this study.

| Sample code | SRA access # | Stage | Part of insect body | Origin | Date of collection | Other information | Refence |
| --- | --- | --- | --- | --- | --- | --- | --- |
| A | SRX3035962 | Larvae | Whole body | Mississippi, USA | March 2016 | Polygynous | Allen et al., (2018) |
| B | SRX3035961 | Larvae | Whole body | Mississippi, USA | March 2016 | Polygynous | Allen et al., (2018) |
| C | SRX3035960 | Larvae | Whole body | Mississippi, USA | March 2016 | Polygynous | Allen et al., (2018) |
| X | SRX3035959 | Pupa | Whole body | Mississippi, USA | March 2016 | Polygynous | Allen et al., (2018) |
| Y | SRX3035964 | Pupa | Whole body | Mississippi, USA | March 2016 | Polygynous | Allen et al., (2018) |
| Z | SRX3035963 | Pupa | Whole body | Mississippi, USA | March 2016 | Polygynous | Allen et al., (2018) |
| Q2 | DRX037806 | Adults | Whole body | Texas, USA | September 2012 | Polygynous, queen | Morandin et al., (2016) |
| W2 | DRX037809 | Adults | Whole body | Texas, USA | September 2012 | Polygynous, worker | Morandin et al., (2016) |
| W3 | DRX037810 | Adults | Whole body | Texas, USA | September 2012 | Polygynous, worker | Morandin et al., (2016) |
| Y05 | SRX5464977 | Adults | Brain | Florida, USA | May 2014 | Queen | - * |
| K05 | SRX5464984 | Adults | Brain | Florida, USA | May 2014 | Queen | - |
| CA01 | SRX5464990 | Adults | Brain | Florida, USA | May 2014 | Queen | - |
| 2-small | SRX5822389 | Adults | Whole body | Taiwan | May 2014 | A pool of two virgin queens, lab colony | Fontana et al., (2020) |

\* Reference is not available yet.

## Methods

### Data set selection and sequencing

Contigs corresponding to a putative new virus genome were initially obtained from a transcriptome project studying differential gene expression between larvae and pupae stages of *S. invicta* collected in Mississippi, U.S. (36). To further investigate and characterize viral diversity associated with *S. invicta*, six libraries (36) were downloaded from GenBank’s Sequence Read Archive (SRA) and analyzed. In addition, to determine whether those viruses were present in other locations in U.S. and abroad, several transcriptome libraries deposited at SRA were compared against those previous putative viral contigs using BLASTn. Libraries that presented high abundance of reads mapping to putative viral contigs with an E-value cutoff lower than 1e-2 were downloaded and further analyzed. The final data set analyzed here consisted of 13 libraries, all from *S. invicta* transcriptomes. Detailed information on the samples are presented in Table 1. All information about sample collection, RNA extraction, library preparation and high throughput sequencing has been previously described (36–38).

### *De novo* assembly and virus genome characterizations

Sequencing reads were trimmed for quality (Suppl. Table S1) and *De novo* assembled using CLC Genomics Workbench 20.0.1 with default settings. Reads associated to *S. invicta* were first filtered by mapping them back to the *S. invicta* genome (GenBank accession: GCA_000188075.2) and only unmapped reads were used for *De novo* assembling (Suppl. Table S1). Contigs were compared against NCBI non-redundant protein sequence (nr) using BLASTx and those predicted to contain near full-length and intact coding sequences homologous to viral sequences previously described were confirmed by mapping reads back to obtain consensus sequences for each library. Potential open reading frames (ORFs) of the putative new viruses were predicted using ORF finder (NCBI) and by comparative analysis with those related viruses. For identification of conserved functional domains, a domain-based BLAST search was performed against the Conserved Domain Database (CDD). ORFs that did not present any conserved domain were predicted based on comparative analysis with other known closely related viruses (5).

### Multiple sequence alignment and phylogenetic analysis

For phylogenetic analyses, one representative sequence for each taxon most closely related to viruses described here was retrieved from GenBank according to BLASTx analysis. ORF integrity was checked and extracted using ORF finder and the presence of conserved domain characteristics for each group was checked at CDD before proceeding to phylogenetic analysis. For highly divergent data sets, regions comprising protein conserved domains were extracted using a script written in R software according to coordinates obtained from CDD for each sequence, and only the conserved domain was used (as indicated below in the figure legends). Multiple sequence alignments of deduced amino acid sequences were prepared using the MUSCLE option in MEGA7 (39). Alignments were manually checked and adjusted when necessary. Phylogenetic trees were constructed using Bayesian inference performed with MrBayes 3.2.6 (40). Estimates of the amino acid model were automatically conducted by setting the prior for the amino acid model to mixed, as implemented in MrBayes. The analyses were carried out running two independent runs of 20,000,000 generations with sampling at every 1000 generations and a burn-in of 25%. Convergence between runs were accepted when average standard deviations of split frequencies was lower than 0.01. Trees were visualized and edited using FigTree and Corel Draw, respectively.

### Inferring viruses-*S. invicta* association

Most samples used here were prepared from whole body ants, and to differentiate truly replicating viruses from those that might be from food contamination or commensal organisms, viral abundance was compared with abundance of three housekeeping genes: cytochrome c oxidase subunit I (*cox1*), that presented high expression levels, ribosomal protein L18 (*rpl18*) and translation elongation factor 1 (*eif1-beta*), both with lower expression levels. Viral and internal genes abundance were calculated as percentage of viral or housekeeping reads per total number of reads in each library (Table 2). Differences among groups was assessed using non-parametric Kruskal-Wallis test followed by post hoc multiple comparison test using Fisher’s least significant difference (41) using Agricolae package (42) in R software. The data were log-transformed before statistical testing. Thus, a virus was suggested to be hosted by *S. invicta* if they qualified in at least two of the criteria as previously suggested by (8) with slight modifications: (i) virus abundance was within the range or higher abundance than housekeeping genes; (ii) viral reads per total number of reads in a library was higher than 0.01%; (iii) they were phylogenetically close to another insect virus; and (iv) complete viral coding sequence regions were recovered.

**Table 2.**
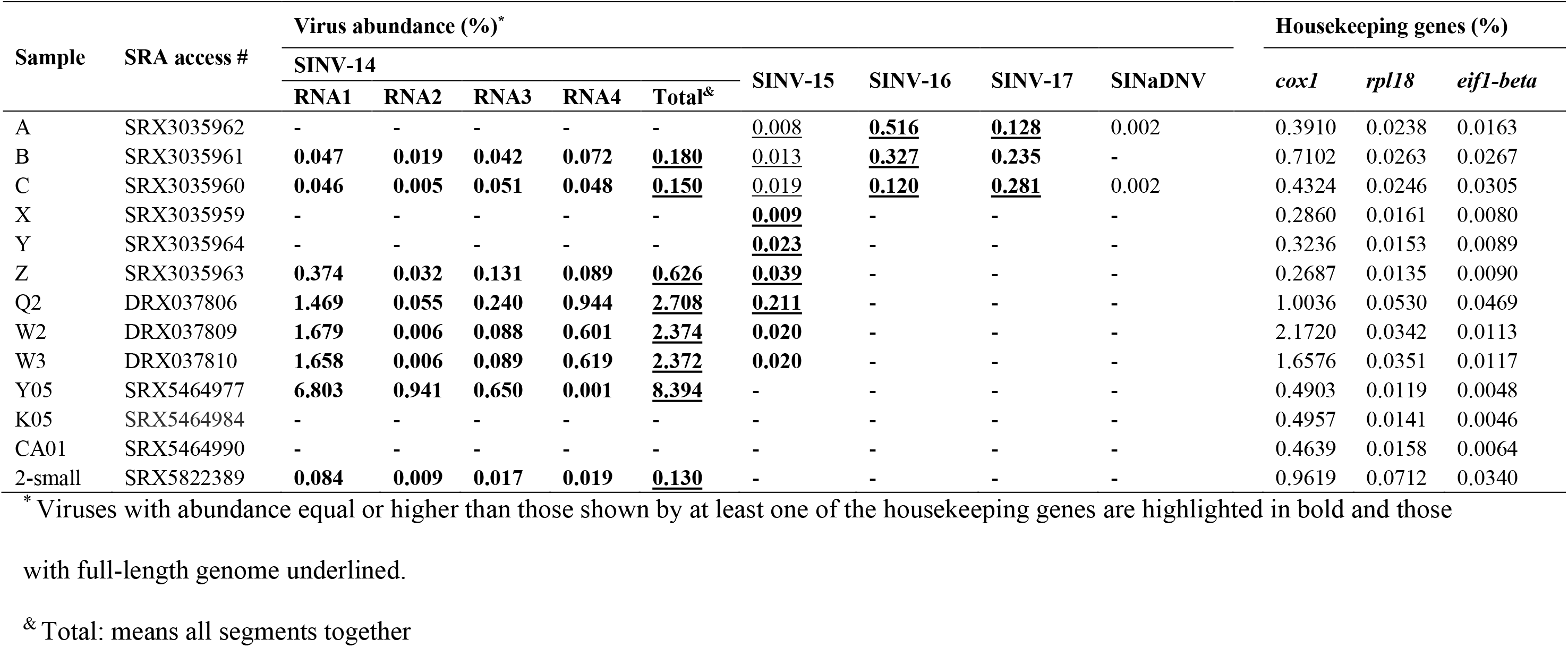
Abundance of viruses and housekeeping genes, cytochrome c oxidase subunit I (*cox1*), ribosomal protein L18 (*rpl18*) and translation elongation factor 1 (*eif1-beta*), expressed in percentage of viral reads per total number of reads in each library.

### Principal component analysis (PCA) of compositional bias

Synonymous codon usage (SCU) and dinucleotide composition bias has been used to infer putative host-virus association and taxonomic placement for a given virus (43, 44). To gain further insight about the host of the tenui-like sequences reported here, we used principal component analysis (PCA) to compare the compositional bias between plant tenuivirus and other viruses infecting plant and animals (vertebrates and invertebrates). SCU counts were determined for the full-length nucleotide sequence of RNA-dependent RNA polymerase (RdRp), glycoprotein, nucleocapsid and non-structural protein 4 (NS4) protein, using the coRdon package (45) in R software. Dinucleotide bias was calculated as the frequency of dinucleotide *XY* divided by the product of frequencies of nucleotide *X* and nucleotide *Y* using the SeqinR package (46) in R software. Sequences highly divergent with no conserved domain according to CDD analysis were not included. Principal components analysis was performed on codon usage counts and dinucleotide bias using the function prcomp and plotted using the factoextra package (47), both implemented in R software.

## Results

### New viruses found associated with *S. invicta* transcriptome

Virus sequences were found associated with *S. invicta* transcriptomes collected from three geographic locations in the U.S. and one location from Taiwan, representing different stages of the *S. invicta* life cycle (Table 1). *De novo* assembling of non-host reads (Suppl. Table S1) from 13 libraries followed by BLAST analyses revealed the presence of complete or near-complete genomes of 4 putative new single-strand RNA (ssRNA) viruses, tentatively named S. invicta virus 14 (SINV-14), SINV-15, SINV-16 and SINV-17, and one partial genome encompassing the almost full-length coding sequence of a ssDNA virus, named S. invicta-associated densovirus (SINaDNV; Figure 1). Sequence comparisons revealed extensive divergence across ORFs and genomic regions with most of them sharing less than 40% amino acid identity compared to those closest homologous sequences previously characterized (Suppl. Table S2). In addition, typical domains associated with RdRp and other proteins involved in virus replication, encapsidation, and entry into the cell were found and characteristic of known virus groups, again confirming the viral identity of those sequences (Suppl. Table S3). Further analyses demonstrated that SINV-14 and SINV-15 fit within the order *Bunyavirales* and *Mononegavirales*, respectively, with SINV-14 having a multisegmented genome (Figure 1). The two other viruses with non-segmented +ssRNA genomes, SINV-16 and SINV-17, were included within the family *Iflaviridae* and into an unclassified group closely related to a proposed new genus of insect-infecting viruses “*Negevirus*” (48), respectively. Finally, the partial ssDNA virus SINaDNV was included into family *Parvoviridae* subfamily *Densovirinae*.

**Figure 1.**
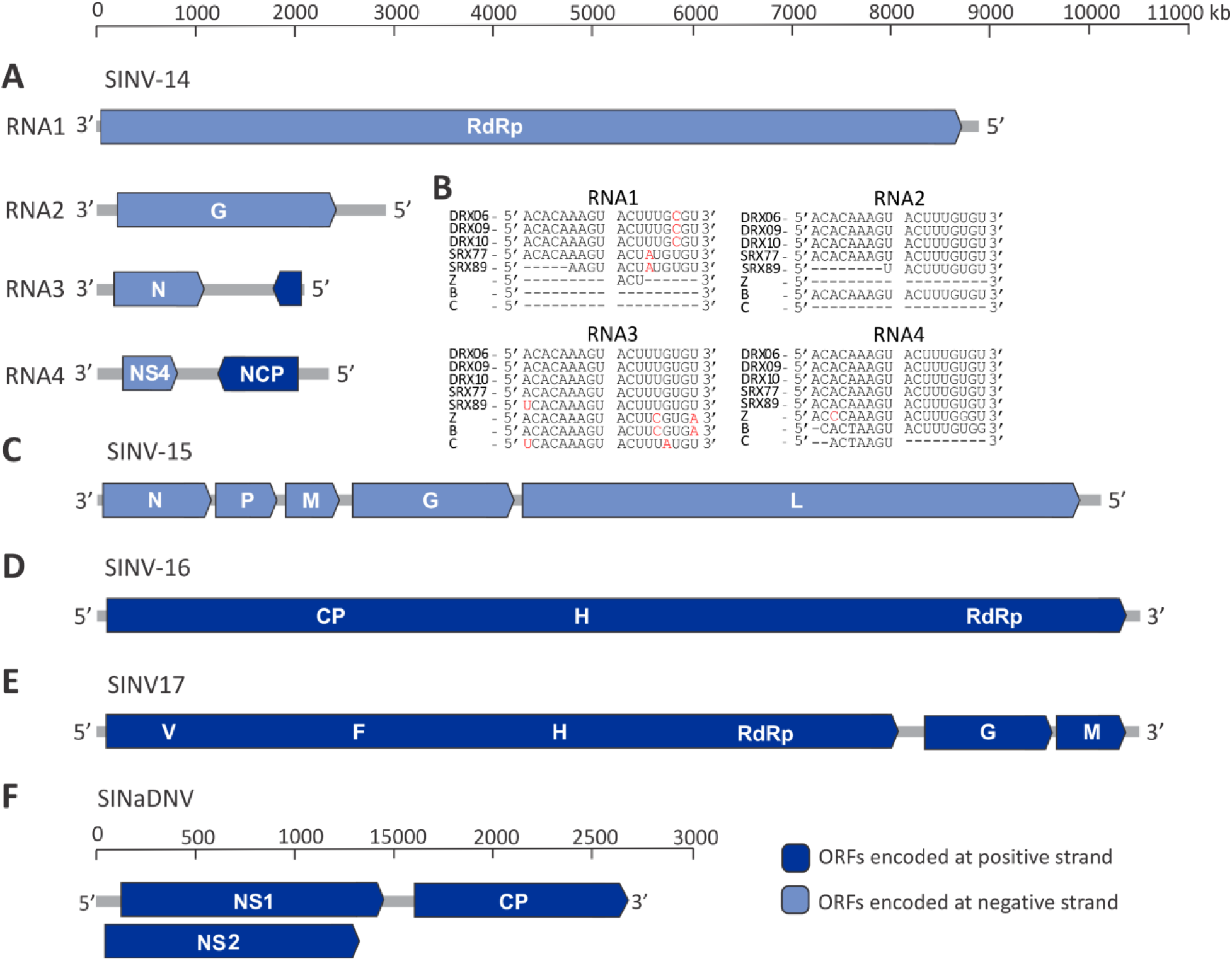
Genome organization of five putative new viruses associated with *S. invicta* transcriptomes. Predicted open reading frames (ORFs) encoding proteins homologous to known viral proteins or conserved domain that they encode are represented and colored according to their orientation in the genome (dark blue: those encoded on positive strand and light blue: those encoded on negative strand). Graphic representation of (**A**) segmented negative single-strand RNA genome S. invicta virus 14 (SINV-14) and (**B**) conserved consensus sequence located at the ends of tenuivirus genome segment (5’-ACACAAAGU and ACUUUGAGU-3’) detected in SINV-14 genome. Dashed lines indicate incomplete sequence. Nucleotides highlighted in red differ from consensus sequence. (**C**) Negative-sense single-strand RNA genome S. invicta virus 15 (SINV-15), and positive-sense single-strand RNA genomes (**D**) S. invicta virus 16 (SINV-16) and (**E**) S. invicta virus 17 (SINV-17) are shown. (**F**) Partial double-strand DNA genome of *S. invicta*-associated densovirus (SINaDNV) is represented. RdRp and L: RNA-dependent RNA polymerase; G: glycoprotein; N: nucleocapsid protein; p3: unknown; NS4: non-structural protein 4; NCP: non-nucleocapsid protein; P: phosphoprotein; M: matrix protein; CP: capsid protein; H: Helicase; V: viral methyltransferase; F: Ftsj-like methyltransferase; NS1and NS2: non-structural protein 1 and 2, respectively.

### Molecular and phylogenetic characterization of new viruses

#### Negative-sense RNA viruses

The SINV-14 genome consists of a linear, negative-sense single-strand RNA composed of four segments, RNA1, RNA2, RNA3 and RNA4 with approximately 8900, 2862, 2086 and 2501 nucleotides (nt), respectively (Figure 1A). Identity analysis of deduced amino acid sequences of coding regions across all segments demonstrated an extensive divergence with those most closely related viruses (Suppl. Table S2). The segments RNA1 and RNA2 encode only one ORF, located on the negative strand (Figure 1A). The deduced amino acid sequence of RNA1 ORF was the most conserved, showing highest identity with the RdRp of Fitzroy Crossing tenui-like virus 1 (FCTenV1), a new tenui-like virus described associated with *Culex annulirostris* from western Australia (Suppl. Table S2) (49), followed by European wheat striate mosaic virus (EWSMV), a typical plant tenuivirus (50). Likewise, even though the RNA2 ORF was the most divergent, it showed highest identity with the glycoprotein of FCTenV1 (Suppl. Table S2). In contrast with RNA1 and RNA2, the segments RNA3 and RNA4 encode two ORFs in ambisense orientation (Figure 1A). Deduced amino acid sequence of RNA3 ORF, encoded on the negative strand, showed highest identity with nucleocapsid protein of otter fecal bunyavirus, an unclassified *Phlebovirus* (51), while no significant similarity was found to a small putative ORF located on the positive strand (Suppl Table S2). The BLASTx analyses indicated that ORFs encoded by RNA4 in the positive and negative strands were related to those of non-nucleocapsid proteins from typical plant tenuiviruses, and nonstructural protein 4 (NS4) of FCTenV1 followed by other plant tenuiviruses, respectively (Suppl. Table S2). Interestingly, the ORF encoded on the negative strand contained the specific domain of movement proteins (pfam03300) of plant tenuiviruses (Suppl. Table S3). A domain-based BLAST search was performed against the CDD, confirming the presence of conserved domains from typical members of family *Phenuiviridae* for RdRp, glycoprotein, putative NS4 and non-nucleocapsid protein (Suppl. Table S3). Also interestingly, further sequence analysis revealed the presence of conserved sequences at the ends of all four segments identical to those found at tenuivirus genomes (3’-ACUUUGUGU and ACACAAAGU-5’; Figure 1B) (52), indicating that full length genome segments were likely assembled from the RNA-Seq datasets.

To further determine the taxonomic placement of SINV-14 within the order *Bunyavirales*, Bayesian phylogenetic trees were inferred based on deduced amino acid sequence of RdRp, glycoprotein, nucleocapsid and NS4, representative of all four segments found (Figure 2). In accordance with identity analysis, SINV-14 RdRp was closest related to that of FCTenV1 (Figure 2A). These two viruses clustered closely to *Horsefly horwuvirus* (virus Wuhan horsefly virus, WhHV), a tenui-like virus described from a poll of horseflies (family *Tabanidae*) (2), the only species within the genus *Horwuvirus* (Figure 2A). Moreover, while the WhHV putative glycoprotein and NS4 were very divergent and no conserved domains were detected based on CDD analyses (thus they were not included into phylogenetic analyses) SINV-14 glycoprotein and NS4 phylogenetic trees were congruent with RdRp, clustering in a well-supported clade closest to FCTenV1 and composing a sister clade with typical plant tenuiviruses (Figure 2B and C). Interestingly, the phylogenetic tree of the nucleocapsid protein was incongruent compared to those of other viral proteins (Figure 2D). It was more closely related to that of otter fecal bunyavirus and other representative viruses of different genus within family *Phenuiviridae* rather than with plant tenuiviruses (Figure 2D). The identity analysis supports the phylogenetic placement in which nucleocapsid protein was significantly closer related to otter fecal bunyavirus than tenui-like and plant tenuiviruses (Suppl. Table S2).

**Figure 2.**
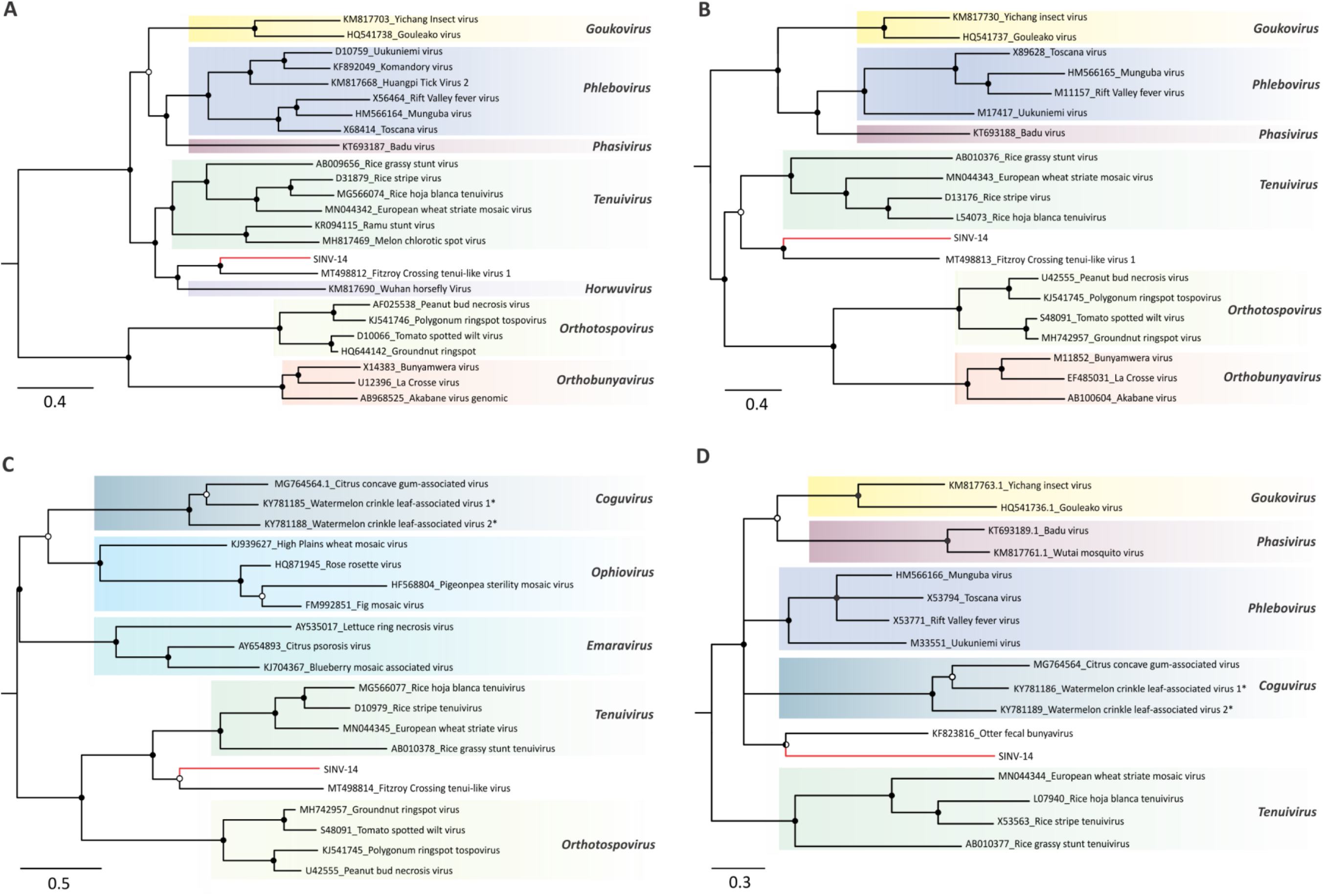
Evolutionary history inferred for different genomic segments of S. invicta virus 14 (SINV-14). Midpoint-rooted Bayesian phylogenetic trees based on (**A**) full-length amino-acid sequences of RNA-dependent RNA polymerase (RdRp), (**B**) amino-acid sequence of conserved glycoprotein domain, (**C**) full-length amino-acid sequence of movement protein and (**D**) full-length amino-acid sequence of nucleocapsid protein. Bayesian posterior probabilities are shown at the nodes. Nodes with posterior probability values between >0.50 and ≤85 are indicated by empty circles, >0.85 and <0.95 are indicated by half-filled circles and those with values ≥0.95 are indicated by filled circles. The scale bar at the bottom indicates the number of amino acid substitutions per site. Branches highlighted in red indicate virus characterized in this study. Genera are indicated at the right side. Proposed species unaccepted yet by International Committee on Taxonomy of Viruses (ICTV) are followed by an asterisk. Proteins sequences encoded by cognates components for a given virus that do not have conserved domain according to Conserved Domain Database (CDD) were not included.

The SINV-15 genome comprises a linear negative-sense, single strand RNA varying from 10113 to 10135 nt (Figure 1C). The genome organization is typical of viruses within order *Mononegavirales*, which contain the conserved core predicted to encode five ORFs: nucleocapsid (N), phosphoprotein (P), matrix protein (M), glycoprotein (G) and RdRp (L) on the negative strand (Figure 1C). Domain-based BLAST search confirmed that ORFs 1 and 5 carry nucleocapsid and RdRp conserved domains, respectively, whereas no conserved domains were identified for ORFs 2, 3 and 4 (Suppl. Table S3). Thus, these ORFs were inferred based on comparative analysis with other known related viruses. Identity analysis indicated SINV-15 is most closely related to Formica fusca virus 1 (FfusV-1; Suppl. Table S2), another virus within order *Mononegavirales* recently described associated with the ant *Formica fusca* (9). Indeed, phylogenetic analysis based on deduced amino acid sequence of RdRp clustered SINV-15 closest to FfusV-1 and Formica exsecta virus 4 (FeV-4), both currently unaccepted species yet. As these three viruses composed a clade closely related to *Orinoco virus* (ONCV), the only accepted species of genus *Orinovirus*, it suggests they are putative new members of this genus within the family *Nyamiviridae*. This is the first report of an orinovirus associated with *S. invicta*.

#### Positive-sense RNA viruses

In addition to SINV-14 and SINV-15, which have negative sense genomes, we also characterized two positive sense RNA viruses (Figure 1D and E). SINV-16 has a positive sense single-stranded RNA genome varying from 10,355 to 10,369 nt (Figure 1D). Genome organization is typical of viruses included in the order *Picornavirales*. The single ORF predicted encodes a large polyprotein (2910 amino acids) including RdRp, helicase, and the canonical picornavirus capsid protein domain (Figure 1D and Suppl. Table S3). Sequence comparison based on amino acid sequence of the polyprotein revealed significant identity with two unaccepted iflavirus-related species, pink bollworm virus 4, isolated from the pink bollworm moth *Pectinophora gossypiella* (53) and Hubei myriapoda virus 1, isolated from a pool of individuals of genus *Scutigeridae* (3). Phylogenetic analysis of RdRp domain clearly clustered SINV-16 within genus *Iflavirus* (Figure 3B). In accordance with identity analysis, it was most closely related to Hubei myriapoda virus 1, in a well-supported clade composed also by pink bollworm virus 4 and *Dinocampus coccinellae paralysis virus* (DcPV), an accepted iflavirus species isolated from the parasitic wasp *Dinocampus coccinellae*. These results indicate that SINV-16 is a new iflavirus reported from *S. invicta*.

**Figure 3.**
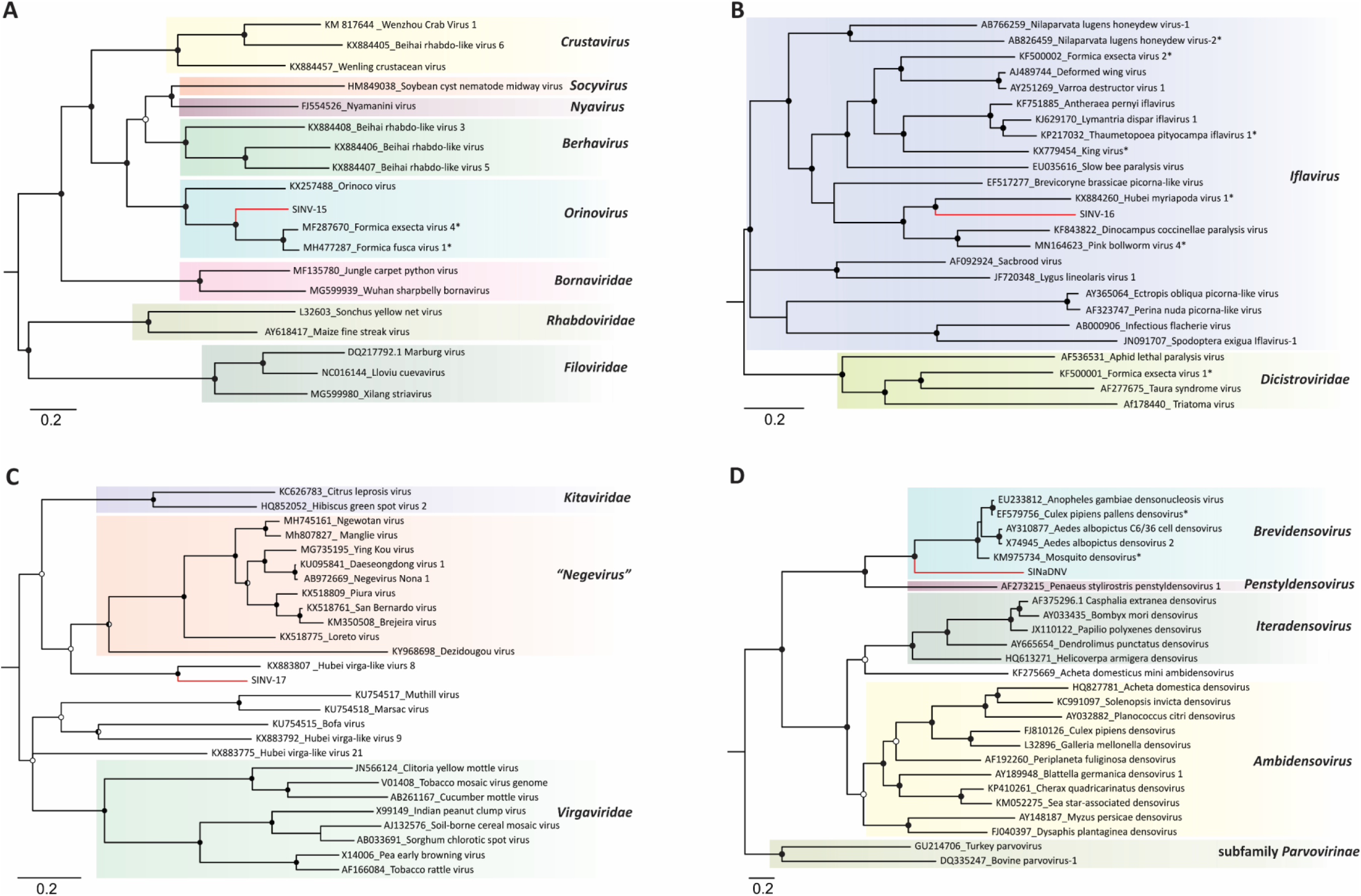
Evolutionary history of new viruses associated to *S. invicta* transcriptome based on Bayesian inference. Midpoint-rooted phylogenetic trees were constructed based on (**A**) conserved domain (pfam00946) of RNA-dependent RNA polymerase (RdRp) of S. invicta virus 15 (SINV-15) and representative species of the order *Mononegavirales*, (**B**) conserved domain (pfam00680) of RdRp of S. invicta virus 16 (SINV-16) and representative species of the families *Iflaviridae* and *Dicistroviridae* and (**C**) full-length amino-acid sequences of RdRp of S. invicta virus 17 (SINV-17) and representative species of the proposed new genus *Negevirus* (unassigned to any family) and families *Virgaviridae* and *Kitaviridae*, are shown. (**D**) Bayesian phylogenetic tree based on full-length amino-acid sequences of non-structural gene 1 (NS1) of S. invicta-associated densovirus (SINaDNV) and representative species of the family *Parvoviridae*. Bayesian posterior probabilities are shown at the nodes. Nodes with posterior probability values between >0.65 and ≤85 are indicated by empty circle, >0.85 and <0.95 are indicated by half-filled circles and those with values ≥0.95 are indicated by filled circles. The scale bar at the bottom indicates the number of amino acid substitutions per site. Branches highlighted in red indicates viruses characterized in this study. Genera/Families/Subfamilies are shown at right side.

The SINV-17 genome consists of a linear, positive-sense single-stranded RNA with approximately 10,341 nt predicted to encode three putative ORFs (Figure 1E). ORF1 was predicted to encode the longest protein containing the viral methyltransferase, helicase and RdRp domains (Figure 1E and Suppl. Table S3). ORF2 and ORF3 were predicted to encode smaller structural proteins, which contain domains associated with virion glycoproteins of insect viruses and virion membrane proteins of plant and insect viruses, respectively (Suppl. Table S3). Interestingly, while ORFs 1 and 2 were most similar to Hubei virga-like virus 8, an unclassified virus isolated from a pool of individuals of the genus *Scutigeridae* (3), ORF 3 showed significant identity to Loreto virus isolated from *Anopheles albimanus*, a member of the proposed genus *Negevirus* (48) (Suppl. Table S2). To further determine the taxonomic placement of SINV-17, Bayesian phylogenetic trees were inferred based on the deduced amino acid sequence of ORF1 RdRp domain. In accordance with identity analysis, SINV-17 was most closely related to Hubei virga-like virus 8, appearing to represent a sister clade to the proposed genus *Negevirus*, and possibly representing a new genus (Figure 3C).

#### Other virus sequences

Two small contigs were found showing significant similarity with RNA virus sequences. The first one has 430 nt in length and exhibited highest identity with a hypothetical protein encoded by Beihai tombus-like virus 6 (41.13% sequence identity; 97% coverage; E-value = 4e-17; GenBank access: APG76145.1), isolated from a pool of marine crustaceans within the genus *Penaeidae* (3). The second contig has 330 nt and showed highest identity with NfV-1 polyprotein (59.49% sequence identity; 93% coverage; E-value = 2e-24; GenBank access: AOC55078.1) from tawny crazy ant *Nylanderia fulva* (54).

In addition to the RNA viruses, we also characterized a partial genome encompassing the almost full-length coding sequence of a single-strand DNA virus. The SINaDNV partial genome assembled from RNA-Seq reads was predicted to encode three ORFs on the positive strand, showing a typical organization of viruses within the family *Parvoviridae* (Figure 1F). The ORFs 1 and 2 were overlapped and were most similar to NS1 and NS2 of mosquito-infecting densoviruses, respectively (Suppl. Table 2). In addition, a third ORF was predicted to encode the capsid protein showing significant identity to Aedes albopictus C6/36 cell densovirus (Suppl. Table S2). Phylogenetic analysis based on deduced amino acid sequence of the NS1, clustered SINaDNV in a well-supported branch most closely related to viruses within subfamily *Densovirinae* and genus *Brevidensovirus* (Figure 3D). While these results strongly suggest SINaDNV is a new densovirus species, further investigation will be needed to reveal its whole genome organization.

#### Inferring host association

Although the transcriptomic approach has been shown to efficiently reveal viral diversity associated with a diverse range of samples, inference about host-virus association it is not always a trivial task. *S. invicta* is a generalist predator/scavenger feeding primarily on other insects and small arthropods, both alive and dead. Thus, it is possible that viruses characterized here could be associated with undigested food or even with *S. invicta*-associated microorganisms, since most samples involved the whole ant body (Table 1). To infer host-virus association and differentiate viruses truly replicating from those that may be a contamination, we calculated the percentage of viral reads for each library and compared with three housekeeping genes, *cox1*, *rpl18* and *eif1-beta* (Figure 4 and Table 2). While the abundance of all three housekeeping genes were relatively stable across different libraries, it differed among them, with *cox1* most abundant, followed by *rpl18* and *eif1-beta* (Figure 4A). Likewise, virus abundance was relatively stable within species, except for SINV-14, which showed a great variation between libraries, varying between 0.130 to 8.394% of viral reads compared to the total reads, considering the virus complete genome (Table 2). For SINV-14 and SINV-16, read abundance was significantly higher than *eif1-beta* and *rpl18*, with no difference compared to *cox1* (Figure 4B). Moreover, SINV-17 abundance was significantly lower than *cox1* and higher than *eif1-beta* and *rpl18*. In contrast, SINV-15 abundance was the lowest one among them, being significantly lower than *coxI*, with no difference compared to *rpl18* and *eif1-beta* (Figure 4B). In addition, SINV-14, SINV-15, SINV-16 and SINV-17 abundances were higher than 0.01% of reads compared to entire library (Table 2). The only exception was SINaDNV, that had a very low abundance compared to housekeeping genes and other viruses (Table 2).

**Figure 4.**
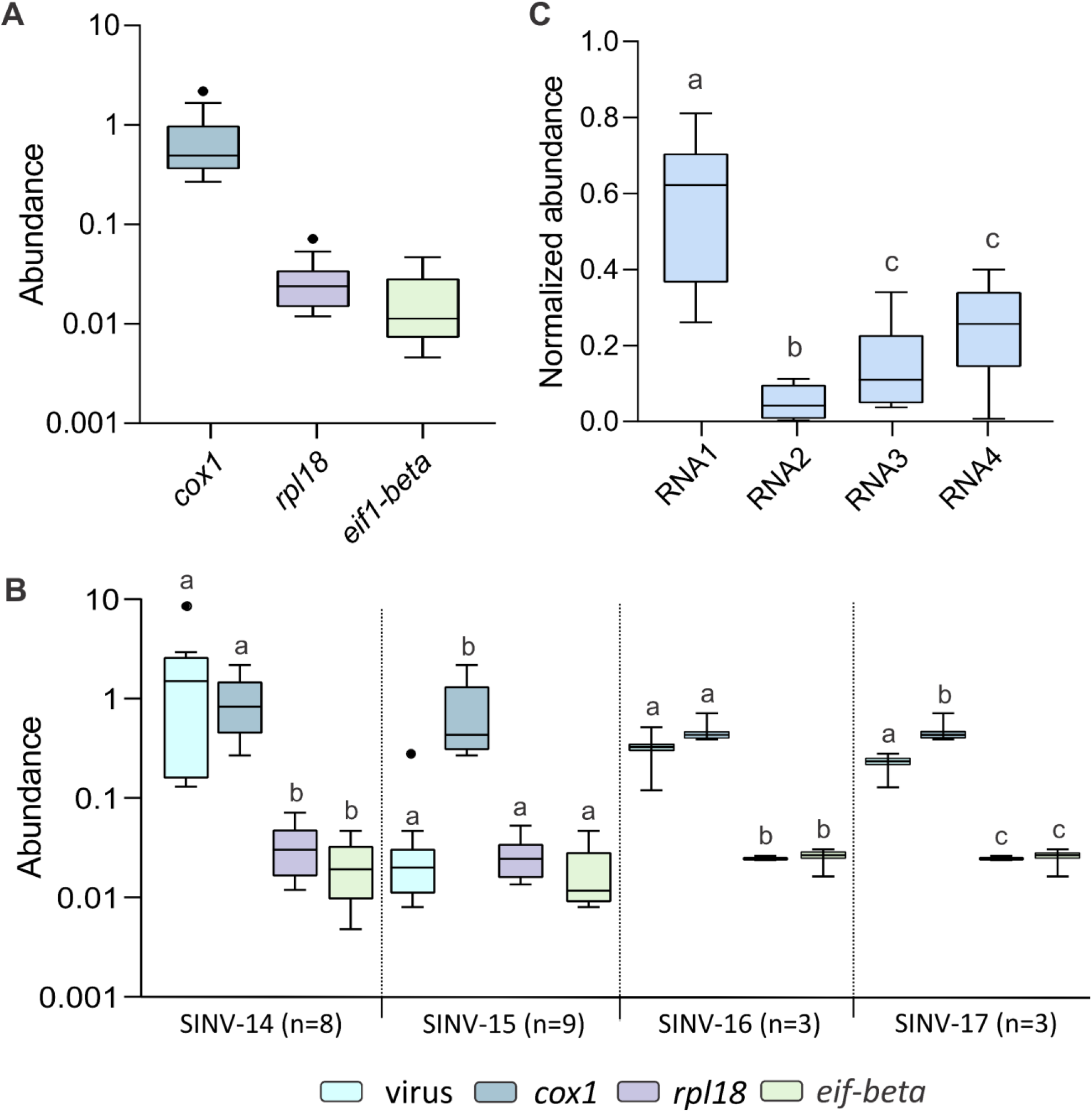
Boxplot of housekeeping genes and viruses abundance. (**A**) Abundance among housekeeping genes from all libraries were compared. Abundance was calculated as percentage of housekeeping or viruses reads per total number of reads in each library. (**B**) Abundance between viruses and housekeeping genes were compared. Statistical test was performed among virus and housekeeping genes abundance from the same libraries. (**C**) Normalized abundance of S. invicta virus 14 (SINV-14) segments significantly differ among them. Normalized abundance was calculated by dividing the abundance of each segment by the total abundance of the entire genome. Box plots with different letters indicate significant differences between groups according to non-parametric Kruskal-Wallis test followed by post hoc multiple comparison test using Fisher’s least significant difference (p<0.05). Box plots show the first and third quartiles as a box, horizontal line corresponds to the median and whiskers correspond to 1.5 times the inter-quartile distance (IQR) from the difference between first and third quartiles.

Interestingly, SINV-14 abundance was much higher in a sample prepared from a dissected brain compared to those from the whole body (Table 1 and 2). Considering the abundance of all segments, SINV-14 from the brain sample was 17-fold more abundant compared with *cox1* gene, whereas abundance of those samples prepared from whole body varied 0.135 to 2.69-fold higher than *cox1* (Table 2). Moreover, the abundance of segments was asymmetric (Figure 4C). The reads mapping on RNA1 were significantly more abundant, followed by RNA3 and RNA4, with RNA2 the least abundant (Figure 4C). These results suggest that SINV-14 is unlikely to be a contamination, and may exhibit a tissue tropism, as we observed evidence that it accumulates more in the brain compared with the whole body.

To further verify the relationship between SINV-14 and *S. invicta*, we also analyzed the codon usage and dinucleotide bias (Figures 5). Compositional biases in virus genomes are likely to mimic those observed in their host species (43, 44), and therefore has been used to assign a given virus to a host and assist in taxonomic placement. Similar data set used for phylogenetic analyses was used here (Suppl. Table S4). Principal component analysis of codon usage based on RdRp, glycoprotein, nucleocapsid and NS4 protein, clearly separate SINV-14 from plant viruses (tenuivirus and orthotospovirus), and non-plant viruses (phenuivirus, orthobunyavirus and tenui-like viruses; Figure 5A-D). Based on the RdRp, three very clear clusters were observed, representing plant virus, non-plant virus and SINV-14 (Figure 5A). Interestingly, the two tenui-like viruses (WhHV and FCTenv1) were closer to plant tenuiviruses than SINV-14, based on RdRp (Figure 5A). Moreover, while SINV-14 clearly clustered separately from other groups, based on glycoprotein, nucleocapsid and NS4, this separation was not very clear between plant and non-plant viruses and tenui-like viruses, as observed for RdRp (Figure 5A-D). We also performed principal component analysis on nucleotide bias obtaining similar results to the codon usage analysis (Figure 5E-H). Whereas RdRp clusters was not very clear, based on plant and non-plant viruses, for all other genes SINV-14 clearly separated from any other group (Figure 5E-H), and again, tenui-like viruses (WhHV and FCTenv1) were closer to other groups than SINV-14.

**Figure 5.**
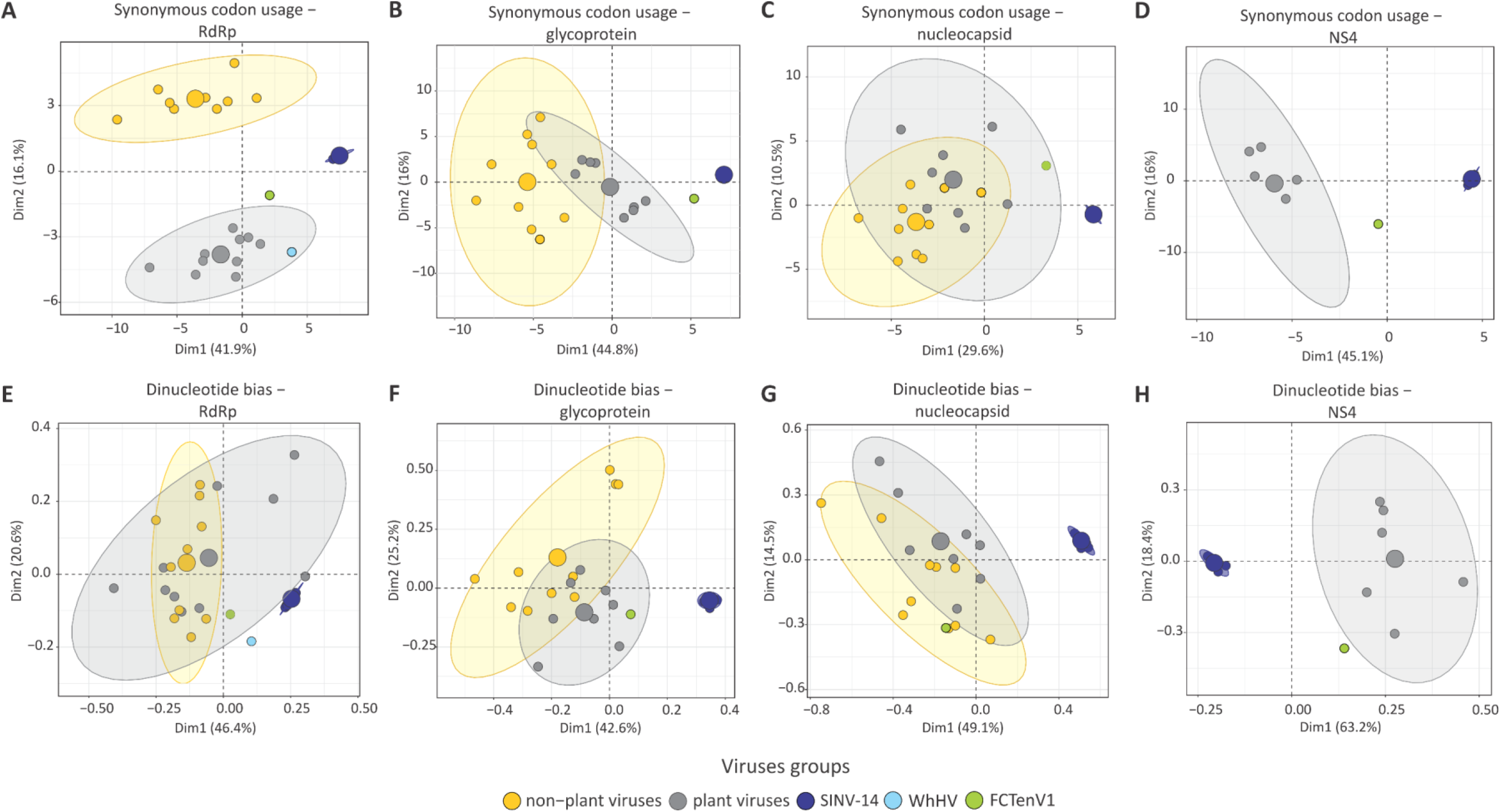
Compositional biases of Solenopsis invicta virus 14 (SINV-14) differ from plant tenuivirus and tenui-like viruses associated with flies. Principal Components Analysis (PCA) of synonymous codon usage (**A**-**D**) and dinucleotide bias (**E**-**H**) of SINV-14 and other viruses that infect plants and animals (vertebrate and invertebrate) are shown. Scatterplots show the first two principal components that account for the largest variance (shown in parenthesis) in the data set with ellipses representing each group and dots individuals. Viruses were divided into two main groups: plant viruses include those that replicate in plants and insect vectors belonging to the genera *Tenuivirus* and *Orthotospovirus* and non-plant viruses including those that do not replicate in plants belonging to the genera *Orthobunyavirus* and representatives within family *Phenuiviridae*. For detailed information of the data set see Supplementary Table S4. For NS4 protein, comparison was performed only for SINV-14, Fitzroy Crossing tenui-like virus1 (FCTenV1) and tenuiviruses, as this gene is not shared by other groups. WhHV, Wuhan horsefly virus.

## Discussion

The advancement of sequencing tools, the collection of numerous genetic samples of high-impact invasive ants, and the continuing discovery of novel insect-related (and other) viruses have been combined to facilitate identification of novel viruses like those reported here. While virus identification for the purpose of addressing pathogenic relationships will certainly continue, it is now possible to identify virome entities that are not necessarily pathogens in the organism samples from which they are identified.

If a species of ant is invasive, are the viruses that persist with that species in its invasive range also invasive species? Definitions of “invasive viruses” would argue that only when a virus causes a disease of obvious impact is the virus itself considered invasive (55). Yet it is easy to speculate that a virus arriving with an invasive ant could prove pathogenic to native ants, insects, and insectivores in the invasive region; perhaps even plants and the human food chain could experience impact. More thorough characterization of the virosphere will enhance our understanding of the roles microbes play and how viruses and hosts interact.

*S. invicta* is a ground-dwelling ant that feeds on a broad diet that may include plant and insect exudates, prey, and decaying matter. Thus, the opportunities for exchange of virus particles from the environment, whether native or novel, are many. The social nature of ant colonies where exchanges of biological fluids among colony members of all castes and life stages must occur also facilitates distribution of virus within the colony. Distinguishing clearly between different host/microbe relationships and forms of virus transmission (horizontal and vertical) will be challenging with invasive ant systems.

Using a transcriptome approach, we searched for RNA viruses in the SRA data collected from *S. invicta.* Curiously, while a great diversity of positive and negative sense viruses has been reported from ants (9, 10) and arthropods (2), *S. invicta* virome has been composed mainly of +ssRNA viruses in the order *Picornavirales*, with no negative sense RNA viruses previously reported. One factor that could be responsible for this bias is the selection of polyadenylated RNA for library preparation, in previous studies (11, 30). The utilization of unbiased library preparation, using ribosomal depletion methods, besides the detection of +ssRNA viruses, has enabled the discovery of many viruses with non-polyadenylated genome, especially within order *Bunyavirales* (2). The fact of some of the libraries used here were prepared using this approach (36) allowed us to characterize SINV-14, the first bunyavirus reported from ants. In addition, we also reported for the first time another negative sense virus within the order *Mononegavirales* associated to *S. invicta*. Although we did not perform any amplification step of viruses genomes characterized here, the presence of untruncated ORFs carrying intact functional domains (Suppl. Table S3), the high abundance of viral reads, and the similar organization and genomes sizes compared to other closely related viruses strongly suggest that we obtained the correct full-length or near full-length genomes (Figure 1). The only exception is SINaDNV, due to the linear single strand DNA (ssDNA) genome. Only active transcriptional units were sampled using our transcriptome approach and further investigation is needed to reveal its full-length genome structure.

Our analysis of RNA obtained from geographically and temporally different samples suggests shifts in the virome of native and invasive fire ants (11). In established exotic populations of the *S. invicta* Yang, Yu (21) identified SINV-1 and SINV-2 and hypothesized that while these two viruses may persist, the more virulent virus, SINV-3 arrived with founders but caused high host mortality resulting in individual carriers of the virus being rapidly eliminated. Nine additional viruses were identified in *S. invicta* RNA samples from the native South American range of the species (11). The viruses described here show distribution and composition that varied according to geographic location and *S. invicta* stage (Table 1 and 2). While SINV-16, SINV-17 and SINaDNV were exclusively found in Mississippi and specifically associated with larvae individuals, detected in all three libraries, these viruses were not detected in pupae from the same sampled place, suggesting a stage-dependent association, as previously reported for some other *S. invicta* viruses (28). Conversely, SINV-14 and SINV-15 were detected associated with all stages analyzed (Table 1 and 2). In addition, SINV-14 and SINV-15 were found in Mississippi and Texas, with SINV-14 also detected in Florida (Table 1 and 2). Interestingly, while SINV-15, SINV-16, SINV-17 and SINaDNV were restricted to the U.S., SINV-14 was also detected in Taiwan (Table 1 and 2). To date, four of five viruses previously reported associated with *S. invicta* in introduced areas have also been previously described to occur in its native origin (11, 21). Although viral diversity associated with *S. invicta* has been deeply studied in Argentina, the viruses reported in this study have not been detected there yet (11, 30). To gain further insight about the new viruses’ distribution in *S. invicta* native range, eight transcriptome libraries publicly available at SRA, were analyzed. These libraries were prepared from 182 colonies collected from eastern Formosa, Argentina, suggested to be the original source to *S. invicta* population in U.S. (11, 18). Based on BLASTn analysis, our search failed to detect any significant reads matching to virus genomes obtained here (data not shown), suggesting that these viruses may not be present in the sampled area in Argentina, and even that new host-virus associations may have occurred in introduced areas.

While prospecting viral enemies in the origin center of *S. invicta* offers opportunity for discovering a higher viral diversity with potential to be used in biocontrol, investigating viruses circulating in introduced populations may reveal new virus-ant interactions with even greater potential. Due to the short-term virus-host association, those viruses acquired in introduced areas may be even more harmful to the host. Furthermore, selecting natural enemies already present in the invaded area is likely to offer less risk to the natural ecosystem. Thus, the viruses characterized here expand the diversity of viruses infecting *S. invicta* in invaded areas with potential to be used as biological control agents, which will require further biological characterization.

The use of metatranscriptome approach has allowed the discovery of divergent groups of RNA viruses contributing to a better understanding about origin and evolutionary history of viruses associated with the most diverse host taxa (56). While the origins of viruses that replicate in plant and insect vectors remains still unknown, such as those within order *Bunyavirales* and *Mononegavirales*, the discovery of possible intermediate forms has been suggested (2, 49). Tenuiviruses are known to be plant viruses that replicate in the insect vector with a multipartite negative/ambisense, ssRNA genome, packaged in a filamentous nonenveloped virion and horizontally transmitted to the host plant in a circulative propagative manner by planthoppers (Order Hemiptera), where vertical transmission can also occur (57, 58). Li, Shi (2) reported the first species of a tenui-like virus, *Horsefly horwuvirus* (virus WhHV), associated with a pool of horseflies (family *Tabanidae*), and proposed that it may represent an transitional form between plant-infecting virus and arthropod-specific viruses. In addition, a partial genome of a new tenui-like, FCTenV1, has been recently reported associated to *Culex annulirostris* (49). While these two tenui-like viruses have been reported associated with flies (Order Diptera), we characterized a new tenui-like virus closely related to the FCTenV1, associated to *S. invicta* (Order Hymenoptera). Interestingly, in contrast with WhHV, which lacks an ambisense coding strategy, the SINV-14 genome exactly mirrors the genomic structure of typical plant tenuivirus, encoding proteins using ambisense strategy, and also has the conserved sequences at the ends of all four segments identical to those found in tenuivirus genomes (Figure 1A and B). While these viruses may represent different steps of transitional forms between tenui-infecting plant and those insects-specific viruses, the direction of the process, if they come from plant to insect or otherwise, remains unknown.

Whereas phylogenetic trees based on RdRp, glycoprotein and NS4 protein were highly congruent and related to those from tenui-like viruses, the nucleocapsid protein was closely related to that of otter fecal phlebovirus, strongly suggesting that SINV-14 may have a recombinant/reassortment origin, where the RNA3 or part of it was acquired from a divergent phenuivirus. While it could be argued that this might be an artefact due to assembling a segmented genome using a transcriptome approach, the fact that we did not find any other contigs related to phenuiviruses, the high read abundance, the constant association between these four segments across different libraries and the presence of the conserved tenuivirus sequence located at the ends of genome segments (Figure 1B), suggest that they are part of a unique genome, rather than being an artefact. In addition, phylogenetic congruence among different segments indicate that they are in an intimate codivergence process (Suppl. Figure S1).

Valles, Strong (59) using an expressed sequence tag (ETS) library from *S. invicta*, detected a short sequence (approximately 750 nt, GenBank access: EF409991) related to plant tenuiviruses. Further identity analysis showed that sequence is 99.8% identical to SINV14 RNA4. They suggested that the sequence would be likely a contamination due to the ant diet feeding either plant or infected insect. Tenuiviruses are typical plant viruses that replicate in the insect vector (57, 58), and the high abundance of SINV-14 compared to housekeeping genes is strong evidence of active replication in *S. invicta*. Furthermore, the highest virus abundance was found in a sample prepared from a dissected ant brain, which rules out the possibility of contamination due to feeding on plants or association with insect vectors infected with a tenuivirus. The Maize stripe virus (MSpV), a typical plant tenuivirus, was detected in the brain of its vector *Peregrinus maidis*, providing evidence of replication of tenuiviruses in this tissue (60). Tenuiviruses replicate in diverse tissues of their insect vectors and are transovarially transmitted between generations suggesting that SINV-14, and other tenui-like, could be sustained through vertical transmission in their insect host (58). Additionally, significant asymmetric abundance of different components of SINV-14 suggest a very specific and active interaction. Asymmetric accumulation in multipartite viruses has been shown and seems to be common trait, shared by RNA and DNA multipartite viruses infecting plants and animals (61, 62); this has been suggested to be involved in control of gene expression allowing fast virus adaptation (63). Although this has not yet been shown for any phenuivirus and may be host dependent (61), our results suggest that this may occur, at least for SINV-14, and more experiments will be necessary to confirm.

While SINV-14 is evolutionarily closely related to insect tenui-like viruses and plant tenuivirus, the synonymous codon usage and dinucleotide analyses demonstrate a distinct compositional bias compared to FCTenV1 and WhHV, and all other viruses analyzed here, indicating that the virus may be actively replicating in ants rather than plants and other insects. The active replication may have driven the virus genome to distinct compositional bias at the nucleotide level, while maintaining protein integrity at the amino acid level and close relationship with those of other tenui-like viruses, as observed through phylogenetic analysis. The fact that most viruses examined here replicate in plant or vertebrate host and also in the insect vector, could be the reason driving such difference between SINV-14, most likely associated only with ant, compared to other viruses. Furthermore, phylogenetic congruence across different segments that mirror the genetic structure of invasive *S. invicta*, suggest an intimate and long-term codivergence process between virus-host. Ascunce, Yang (18) demonstrated that *S. invicta* subpopulations from Texas comprise a distinct genetic cluster compared to Mississippi and Florida, that can also be observed when we reconstructed the phylogeny for the distinct segments of SINV-14 (Suppl. Figure S1). Interestingly, while Ascunce, Yang (18) suggested an introduction event from California into Taiwan, the virus recovery from Taiwan clustered with those from Mississippi and Florida, suggesting that other more recent introductions may have occurred (64, 65).

We presented strong evidence that SINV-14 is long-term association and may actively replicate in ants. However, the possibility of this virus, and other tenui-like to replicate in plants is unknown. The presence of the protein carrying an NS4 domain, that is related to cell-to-cell and long-distance movement in plants (66) in insect viruses is puzzling. Solenopsis invicta virus 14 NS4 is highly divergent showing 19.85 to 24.4% of identity compared to other tenuivirus (Suppl. Figure S2). In addition, mutation in most sites associated with cell-to-cell and long-distance movement (Suppl. Figure S2), suggests that this protein might have lost the capacity to move viral genome in plant, whereas its maintenance in insect viruses may be related to another role acquired through functional diversification. The possible function of the plant virus movement protein from insect viruses is of significant interest, and its role in insects and plants remains to be addressed.

Altogether, based on virus abundance compared to housekeeping genes, abundant viruses with viral reads higher than 0.01%, phylogenetic relationship, complete viral coding sequence regions recovered, and compositional bias for SINV-14, our results suggest that four out five viruses reported here, those being SINV-14, SINV-15, SINV-16 and SINV-17 are truly replicating in *S. invicta*. Our results suggest fluid shifts in the virome of this invasive species. Further research describing this virome in native and invasive regions and ecosystems could provide insight on virus evolution and invasion mechanics.

## Conclusions

The present study expands our knowledge about viral diversity associated with *S. invicta* in introduced areas. In addition to revealing new virus-host interactions, contributing to better understanding of viral evolutionary history and emergence, the discovery of new viruses expands the range of agents with potential to be used in biocontrol programs, which will require further biological characterization. By understanding these interactions, we may be better equipped to cope with ongoing global changes and introductions of invasive organisms and their associated viruses.

## Availability of data and materials

Consensus virus genome sequences were deposited in GenBank under accession numbers: MT860232 to MT860267.

## Ethics approval and consent to participate

Not applicable.

## Consent for publication

Not applicable.

## Authors’ contributions

C.A.D.X, M.L.A. and A.E.W. conceived and designed the study. C.A.D.X and M.L.A. performed the analysis. C.A.D.X, M.L.A. and A.E.W. discussed the results and wrote the manuscript.

## Acknowledgements

Trade, firm, or corporation names in this publication are for the information and convenience of the reader. Such use does not constitute an official endorsement or approval by the U.S. Department of Agriculture of the Agricultural Research Service (USDA-ARS).

## References

1. Stork NE. How many species of insects and other terrestrial arthropods are there on Earth? Annu Rev Entomol. 2018;63:31–45.

2. Li CX, Shi M, Tian JH, Lin XD, Kang YJ, Chen LJ, et al. Unprecedented genomic diversity of RNA viruses in arthropods reveals the ancestry of negative-sense RNA viruses. Elife. 2015;29(4):05378.

3. Shi M, Lin X-D, Tian J-H, Chen L-J, Chen X, Li C-X, et al. Redefining the invertebrate RNA virosphere. Nature. 2016;540(7634):539–43.

4. Käfer S, Paraskevopoulou S, Zirkel F, Wieseke N, Donath A, Petersen M, et al. Re-assessing the diversity of negative strand RNA viruses in insects. PLoS Pathog. 2019;15(12).

5. Shi M, Neville P, Nicholson J, Eden JS, Imrie A, Holmes EC. High-resolution metatranscriptomics reveals the ecological dynamics of mosquito-associated RNA viruses in western australia. J Virol. 2017;91(17):00680–17.

6. Atoni E, Zhao L, Karungu S, Obanda V, Agwanda B, Xia H, et al. The discovery and global distribution of novel mosquito-associated viruses in the last decade (2007–2017). Rev Med Virol. 2019;29(6):13.

7. Faizah AN, Kobayashi D, Isawa H, Amoa-Bosompem M, Murota K, Higa Y, et al. Deciphering the virome of *Culex vishnui* subgroup mosquitoes, the major vectors of japanese encephalitis, in Japan. Viruses. 2020;12(3).

8. Pettersson JH, Shi M, Eden JS, Holmes EC, Hesson JC. Meta-transcriptomic comparison of the RNA viromes of the mosquito vectors *Culex pipiens* and *Culex torrentium* in Northern Europe. Viruses. 2019;11(11).

9. Kleanthous E, Olendraite I, Lukhovitskaya NI, Firth AE. Discovery of three RNA viruses using ant transcriptomic datasets. Archives of virology. 2019;164(2):643–7. Epub 2018/11/10.

10. Dhaygude K, Johansson H, Kulmuni J, Sundström L. Genome organization and molecular characterization of the three Formica exsecta viruses-FeV1, FeV2 and FeV4. PeerJ. 2019;20(6).

11. Valles SM, Rivers AR. Nine new RNA viruses associated with the fire ant *Solenopsis invicta* from its native range. Virus Genes. 2019;55(3):368–80.

12. Lacey LA, Grzywacz D, Shapiro-Ilan DI, Frutos R, Brownbridge M, Goettel MS. Insect pathogens as biological control agents: back to the future. Journal of Invertebrate Pathology. 2015;132:1–41.

13. Chen X, Gonçalves MA. Engineered viruses as genome editing devices. Mol Ther. 2016;24(3):447–57.

14. Rode NO, Estoup A, Bourguet D, Courtier-Orgogozo V, Débarre F. Population management using gene drive: molecular design, models of spread dynamics and assessment of ecological risks. Conservation Genetics. 2019;20(4):671–90.

15. Porter SD, Savignano DA. Invasion of polygyne fire ants decimates native ants and disrupts arthropod community. Ecology. 1990;71(6):2095–106.

16. Pimentel D, Zuniga R, Morrison D. Update on the environmental and economic costs associated with alien-invasive species in the United States. Ecological Economics. 2005;52(3):273–88.

17. Callcott A-MA, Collins HL. Invasion and range expansion of imported fire ants (Hymenoptera: Formicidae) in North America from 1918-1995. The Florida Entomologist. 1996;79(2):240–51.

18. Ascunce MS, Yang C-C, Oakey J, Calcaterra L, Wu W-J, Shih C-J, et al. Global invasion history of the fire ant *Solenopsis invicta*. Science. 2011;331(6020):1066–8.

19. Shoemaker DD, Ahrens M, Sheill L, Mescher M, Keller L, Ross KG. Distribution and prevalence of *Wolbachia* Infections in native populations of the fire ant *Solenopsis invicta* (Hymenoptera: Formicidae). Environmental Entomology. 2003;32(6):1329–36.

20. Bouwma AM, Ahrens ME, DeHeer CJ, DeWayne Shoemaker D. Distribution and prevalence of *Wolbachia* in introduced populations of the fire ant *Solenopsis invicta*. Insect Mol Biol. 2006;15(1):89–93.

21. Yang C-C, Yu Y-C, Valles SM, Oi DH, Chen Y-C, Shoemaker D, et al. Loss of microbial (pathogen) infections associated with recent invasions of the red imported fire ant *Solenopsis invicta*. Biological Invasions. 2010;12(9):3307–18.

22. Porter SD, Fowler HG, Mackay WP. Fire ant mound densities in the United States and Brazil (Hymenoptera: Formicidae). Journal of Economic Entomology. 1992;85(4):1154–61.

23. Morrison LW, Porter SD, Daniels E, Korzukhin MD. Potential global range expansion of the invasive fire ant, *Solenopsis invicta*. Biological Invasions. 2004;6(2):183–91.

24. Drees BM, Calixto AA, Nester PR. Integrated pest management concepts for red imported fire ants *Solenopsis invicta* (Hymenoptera: Formicidae). Insect Science. 2013;20(4):429–38.

25. Valles SM, Porter SD, Choi M-Y, Oi DH. Successful transmission of Solenopsis invicta virus 3 to *Solenopsis invicta* fire ant colonies in oil, sugar, and cricket bait formulations. Journal of Invertebrate Pathology. 2013;113(3):198–204.

26. Oi D, Valles S, Porter S, Cavanaugh C, White G, Henke J. Introduction of fire ant biological control agents into the Coachella Valley of California. Florida Entomologist. 2019;102(1):284–6, 3.

27. Valles SM, Strong CA, Dang PM, Hunter WB, Pereira RM, Oi DH, et al. A picorna-like virus from the red imported fire ant, *Solenopsis invicta*: initial discovery, genome sequence, and characterization. Virology. 2004;328(1):151–7.

28. Valles SM, Hashimoto Y. Isolation and characterization of Solenopsis invicta virus 3, a new positive-strand RNA virus infecting the red imported fire ant, *Solenopsis invicta*. Virology. 2009;388(2):354–61.

29. Valles SM, Strong CA, Hashimoto Y. A new positive-strand RNA virus with unique genome characteristics from the red imported fire ant, *Solenopsis invicta*. Virology. 2007;365(2):457–63.

30. Valles SM, Porter SD, Calcaterra LA. Prospecting for viral natural enemies of the fire ant *Solenopsis invicta* in Argentina. PLoS One. 2018;13(2).

31. Allen ML. Near-complete genome sequences of new strain of *Nylanderia Fulva Virus 1* from *Solenopsis invicta*. Microbiol Resour Announc. 2020;9(15):00798–19.

32. Valles SM, Shoemaker D, Wurm Y, Strong CA, Varone L, Becnel JJ, et al. Discovery and molecular characterization of an ambisense densovirus from South American populations of *Solenopsis invicta*. Biological Control. 2013;67(3):431–9.

33. Manfredini F, Shoemaker D, Grozinger CM. Dynamic changes in host-virus interactions associated with colony founding and social environment in fire ant queens (*Solenopsis invicta*). Ecol Evol. 2015;6(1):233–44.

34. Hsu H-W, Chiu M-C, Shoemaker D, Yang C-CS. Viral infections in fire ants lead to reduced foraging activity and dietary changes. Scientific Reports. 2018;8(1):13498.

35. Valles SM. Positive-strand RNA viruses infecting the red imported fire ant, *Solenopsis invicta*. Psyche. 2012;2012:821591.

36. Allen ML, Rhoades JH, Sparks ME, Grodowitz MJ. Differential gene expression in red imported fire ant (*Solenopsis invicta*) (Hymenoptera: Formicidae) larval and pupal stages. Insects. 2018;9(4).

37. Morandin C, Tin MMY, Abril S, Gómez C, Pontieri L, Schiøtt M, et al. Comparative transcriptomics reveals the conserved building blocks involved in parallel evolution of diverse phenotypic traits in ants. Genome Biology. 2016;17(1):43.

38. Fontana S, Chang N-C, Chang T, Lee C-C, Dang V-D, Wang J. The fire ant social supergene is characterized by extensive gene and transposable element copy number variation. Molecular Ecology. 2020;29(1):105–20.

39. Kumar S, Stecher G, Tamura K. MEGA7: molecular evolutionary genetics analysis version 7.0 for bigger datasets. Molecular Biology and Evolution. 2016;33(7):1870–4.

40. Ronquist F, Teslenko M, van der Mark P, Ayres DL, Darling A, Höhna S, et al. MrBayes 3.2: efficient Bayesian phylogenetic inference and model choice across a large model space. Systematic biology. 2012;61(3):539–42. Epub 2012/02/22.

41. Bewick V, Cheek L, Ball J. Statistics review 10: further nonparametric methods. Critical care (London, England). 2004;8(3):196–9. Epub 2004/04/16.

42. De Mendiburu F, Yaseen M. Statistical procedures for agricultural research. R package version 140. https://myaseen208.github.io/agricolae/2020.

43. Di Giallonardo F, Schlub TE, Shi M, Holmes EC. Dinucleotide composition in animal RNA viruses is shaped more by virus family than by host species. Journal of virology. 2017;91(8):e02381–16.

44. Kapoor A, Simmonds P, Lipkin WI, Zaidi S, Delwart E. Use of nucleotide composition analysis to infer hosts for three novel picorna-like viruses. J Virol. 2010;84(19):10322–8.

45. Elek A, Kuzman M, Vlahovicek K. coRdon: codon usage analysis and prediction of gene expressivity. R package version 170. https://github.com/BioinfoHR/coRdon2020.

46. Charif D, Lobry JR. SeqinR 1.0-2: a contributed package to the R project for statistical computing devoted to biological sequences retrieval and analysis. In: Bastolla U, Porto M, Roman HE, Vendruscolo M, editors. Structural Approaches to Sequence Evolution: Molecules, Networks, Populations. Berlin, Heidelberg: Springer Berlin Heidelberg; 2007. p. 207–32.

47. Kassambara A, Mundt F. factoextra: extract and visualize the results of multivariate data analyses. R package version 107. http://www.sthda.com/english/rpkgs/factoextra2020.

48. Vasilakis N, Forrester NL, Palacios G, Nasar F, Savji N, Rossi SL, et al. Negevirus: a proposed new taxon of insect-specific viruses with wide geographic distribution. Journal of virology. 2013;87(5):2475–88. Epub 2012/12/19.

49. Williams SH, Levy A, Yates RA, Somaweera N, Neville PJ, Nicholson J, et al. The diversity and distribution of viruses associated with *Culex annulirostris* mosquitoes from the Kimberley region of western Australia. Viruses. 2020;12(7):717.

50. Sõmera M, Kvarnheden A, Desbiez C, Blystad DR, Sooväli P, Kundu JK, et al. Sixty years after the first description: genome sequence and biological characterization of European wheat striate mosaic virus infecting cereal crops. Phytopathology. 2020;110(1):68–79.

51. Bodewes R, Ruiz-Gonzalez A, Schapendonk CME, van den Brand JMA, Osterhaus ADME, Smits SL. Viral metagenomic analysis of feces of wild small carnivores. Virology journal. 2014;11:89-.

52. Toriyama S, Kimishima T, Takahashi M, Shimizu T, Minaka N, Akutsu K. The complete nucleotide sequence of the rice grassy stunt virus genome and genomic comparisons with viruses of the genus *Tenuivirus*. Journal of General Virology. 1998;79(8):2051–8.

53. Dou X. Transcriptome analysis and identification of genes involved in moth sex pheromone biosynthetic pathways. Graduate Theses and Dissertations2019.

54. Valles SM, Oi DH, Becnel JJ, Wetterer JK, LaPolla JS, Firth AE. Isolation and characterization of Nylanderia fulva virus 1, a positive-sense, single-stranded RNA virus infecting the tawny crazy ant, *Nylanderia fulva*. Virology. 2016;496:244–54. Epub 2016/06/30.

55. Invasive.org: center for invasive species and ecossystem healthy. https://www.invasive.org/species/diseases.cfm. Accessed 4 August 2020.

56. Wolf YI, Kazlauskas D, Iranzo J, Lucía-Sanz A, Kuhn JH, Krupovic M, et al. Origins and evolution of the global RNA virome. mBio. 2018;9(6):e02329–18.

57. Falk BW, Tsai JH. Biology and molecular biology of viruses in the genus *Tenuivirus*. Annual review of phytopathology. 1998;36:139–63.

58. Liu W, Hajano JU, Wang X. New insights on the transmission mechanism of tenuiviruses by their vector insects. Curr Opin Virol. 2018;33:13–7.

59. Valles SM, Strong CA, Hunter WB, Dang PM, Pereira RM, Oi DH, et al. Expressed sequence tags from the red imported fire ant, *Solenopsis invicta*: annotation and utilization for discovery of viruses. J Invertebr Pathol. 2008;99(1):74–81.

60. Nault LR, Gordon DT. Multiplication of Maize Stripe Virus in Peregrinus maidis. Phytopathology. 1988;78(7):991–5.

61. Sicard A, Yvon M, Timchenko T, Gronenborn B, Michalakis Y, Gutierrez S, et al. Gene copy number is differentially regulated in a multipartite virus. Nat Commun. 2013;4(2248).

62. Wu B, Zwart MP, Sánchez-Navarro JA, Elena SF. Within-host evolution of segments ratio for the tripartite genome of Alfalfa mosaic virus. Scientific Reports. 2017;7(1):5004.

63. Zwart MP, Elena SF. Modeling multipartite virus evolution: the genome formula facilitates rapid adaptation to heterogeneous environments. Virus Evol. 2020;6(1).

64. Schoebel CN, Botella L, Lygis V, Rigling D. Population genetic analysis of a parasitic mycovirus to infer the invasion history of its fungal host. Molecular Ecology. 2017;26(9):2482–97.

65. Carver S, Lunn T. When are pathogen dynamics likely to reflect host population genetic structure? Molecular Ecology. 2020;29(5):859–61.

66. Zhang C, Pei X, Wang Z, Jia S, Guo S, Zhang Y, et al. The *Rice stripe virus* pc4 functions in movement and foliar necrosis expression in *Nicotiana benthamiana*. Virology. 2012;425(2):113–21.

