## Supplementary information for "Ever-increasing viral diversity associated with the red imported fire ant *Solenopsis invicta* (Formicidae: Hymenoptera)"

**Supplementary Table S1.** Descriptive statistics of fire ant transcriptome datasets analyzed in this study.

| **Sample code** | **SRA access #** | **Total number of reads** | **Number of reads after quality trim** | **Number of reads mapped to *S. invicta* genome** | **Unmapped reads** | **Average length of reads** |
|---|---|---|---|---|---|---|
| A | SRX3035962 | 104,912,554 | 103,820,346 | 99,200,909 | 4,619,437 | 85 |
| B | SRX3035961 | 99,506,994 | 98,441,192 | 95,468,116 | 2,973,076 | 85 |
| C | SRX3035960 | 84,803,184 | 84,400,181 | 61,952,078 | 22,448,103 | 86 |
| X | SRX3035959 | 109,126,076 | 108,107,491 | 105,996,684 | 2,110,807 | 80 |
| Y | SRX3035964 | 117,979,272 | 103,771,205 | 93,478,311 | 10,292,894 | 85 |
| Z | SRX3035963 | 77,554,994 | 77,239,893 | 75,495,885 | 1,744,008 | 85 |
| Q2 | DRX037806 | 36,219,686 | 35,765,978 | 34,263,934 | 1,955,752 | 70 |
| W2 | DRX037809 | 23,526,030 | 23,768,908 | 22,097,772 | 1,428,258 | 77 |
| W3 | DRX037810 | 27,718,678 | 26,345,675 | 26,068,916 | 1,649,762 | 78.01 |
| Y05 | SRX5464977 | 42,874,000 | 41,904,267 | 38,419,996 | 4,454,004 | 125 |
| K05 | SRX5464984 | 41,970,844 | 41,923,483 | 41,243,134 | 680,349 | 105.26 |
| CA01 | SRX5464990 | 52,734,546 | 52,696,226 | 52,097,923 | 598,303 | 105.59 |
| 2-small | SRX5822389 | 42,942,626 | 42,877,252 | 41,671,042 | 1,206,210 | 110.92 |

**Supplementary Table S2.** Open reading frames (ORFs) characterization from putative new viruses based on ORFinder followed by BLASTp analyses, as implemented at National Center for Biotechnology Information Search database (NCBI).

| Group | Segments # | ORF | Strand | Frame | Start | Stop | Length (nt/aa) | Description | Max. score | Total score | Cover | E-value | Identity (%) | Access # |
|---|---|---|---|---|---|---|---|---|---|---|---|---|---|---|
| SINV-14 | RNA1 | 1 | - | 2 | 8895 | 166 | 8730/2909 | RNA-dependent RNA polymerase [Fitzroy Crossing tenui-like virus 1] | 1658 | 1658 | 76% | 0.0E+00 | 42.50% | QLJ83469.1 |
| | RNA2 | 1 | - | 2 | 2861 | 465 | 2217/738 | glycoprotein [Fitzroy Crossing tenui-like virus 1] | 164 | 164 | 75% | 3e-38 | 25.44% | QLJ83470.1 |
| | RNA3 | 1 | + | 3 | 39 | 278 | 240/79 | - | - | - | - | - | - | - |
| | | 2 | - | 2 | 1929 | 1009 | 921/306 | nucleocapsid protein [Otter fecal bunyavirus] | 69.3 | 69.3 | 72% | 6.0E-10 | 26.13% | AIB06815.1 |
| | RNA4 | 1 | + | 3 | 2362 | 1772 | 591/196 | non-capsid protein [Urochloa hoja blanca tenuivirus] | 68.6 | 68.6 | 86% | 1.0E-10 | 26.47% | YP009507913.1 |
| | | 2 | - | 2 | 566 | 1420 | 855/284 | nonstructural protein 4 [Fitzroy Crossing tenui-like virus 1] | 102 | 102 | 67% | 1.0E-21 | 27.60% | QLJ83471.1 |
| SINV-15 | 1 | 1 | + | 3 | 69 | 1124 | 1056/351 | nucleoprotein [Formica fusca virus 1] | 234 | 234 | 92% | 9.0E-71 | 42.07% | AYW51534.1 |
| | | 2 | + | 2 | 1205 | 1828 | 624/207 | -* | - | - | - | - | - | - |
| | | 3 | + | 1 | 1924 | 2457 | 534/177 | ORF3 [Formica exsecta virus 4] | 103 | 103 | 98% | 1.0E-24 | 35.43% | AWI42883.1 |
| | | 4 | + | 3 | 2580 | 4211 | 1632/543 | ORF2 [Formica exsecta virus 4] | 332 | 332 | 95% | 4.0E-104 | 34.87% | AWI42882.1 |
| | | 5 | + | 1 | 4297 | 9924 | 5628/1875 | L polymerase RdRp [Formica fusca virus 1] | 2091 | 2091 | 98% | 0.0E+00 | 54.26% | AYW51538.1 |

| | | | | | | | | | | | | | | |
|---|---|---|---|---|---|---|---|---|---|---|---|---|---|---|
| SINV-16 | 1 | 1 | + | 2 | 1088 | 10231 | 9144/3047 | polyprotein [Pink bollworm virus 4] | 514 | 514 | 76% | 1.0E-142 | 36.33% | QID77679.1 |
| SINV-17 | 1 | 1 | + | 2 | 92 | 7939 | 7848/2615 | RdRp [Hubei virga-like virus 8] | 1699 | 1699 | 99% | 0.0E+00 | 37.49% | APG77744.1 |
| | | 2 | + | 3 | 8205 | 9482 | 1278/425 | hypothetical protein [Hubei virga-like virus 8] | 167 | 167 | 83% | 8.0E-44 | 31.81% | APG77746.1 |
| | | 3 | + | 2 | 9497 | 1021 | 705/234 | hypothetical protein 3 [Loreto virus] | 55.8 | 55.8 | 95% | 1.0E-05 | 23.56% | YP009351837.1 |
| SINaDNV | 1 | 1 | + | 1 | 46 | 1482 | 1437/478 | non-structure protein 1 [Mosquito densovirus HB-3] | 184 | 184 | 80% | 8.0E-47 | 33.50% | AJI02597.1 |
| | | 2 | + | 2 | 185 | 1357 | 1173/390 | non-structure protein 2 [Aedes albopictus densovirus 2] | 203 | 203 | 94% | 4.0E-58 | 40.75% | NP694828.1 |
| | | 3 | + | 3 | 1629 | 2696 | 1068/355 | capsid protein [Aedes albopictus C6/36 cell densovirus] | 220 | 220 | 92% | 5.0E-65 | 38.71% | AAM28945.2 |

* No similarity found.

**Supplementary Table S3.** Viral conserved domains detected in ORFs from putative new viruses based on Conserved Domains Database (CDD).

| Group | ORF | Domain | Pfam accession | Description | Interval | E-value |
|---|---|---|---|---|---|---|
| SINV-14 | RNA1 | Bunya_RdRp | pfam04196 | Bunyavirus RNA dependent RNA polymerase; | 1093-1808 | 8.05E-120 |
| | | L_protein_N | pfam15518 | L protein N-terminus. This endonuclease domain is found at the N-terminus of many bunyavirus | 520-586 | 4.15E-09 |
| | | DUF3770 | pfam12603 | Protein of unknown function (DUF3770); This domain family is found in viruses | 648-909 | 1.81E-05 |
| | RNA2 | Tenui_PVC2 | pfam06656 | Tenuivirus PVC2 protein; This family consists of several Tenuivirus PVC2 proteins from | 141-519 | 1.45E-08 |
| | RNA3 | Tenui_N | pfam05733 | Tenuivirus/Phlebovirus nucleocapsid protein. This family consists of several Tenuivirus | 23-207 | 1.42E-11 |
| | RNA4-1 | Tenui_NCP | pfam04876 | Tenuivirus major non-capsid protein; | 20-175 | 1.48E-13 |
| | RNA4-2 | Tenui_NS4 | pfam03300 | Tenuivirus non-structural, movement protein NS4; | 33-219 | 8.74E-16 |
| SINV-15 | 1 | BDV_P40 | pfam06407 | Borna disease virus P40 protein. This family consists of several Borna disease virus P40 | 83-306 | 9.33E-35 |
| | 2 | -* | - | - | - | - |
| | 3 | - | - | - | - | - |
| | 4 | - | - | - | - | - |
| | 5 | Mononeg_RNA_pol | pfam00946 | Mononegavirales RNA dependent RNA polymerase. | 143-994 | 3.35E-92 |
| | | Mononeg_mRNAcap | pfam14318 | Mononegavirales mRNA-capping region V. This V domain of L RNA-polymerase carries a new motif | 1009-1206 | 2.77E-23 |
| | | paramyx_RNAcap | TIGR04198 | mRNA capping enzyme, paramyxovirus family; This model represents a common C-terminal region | 1106-1335 | 5.61E-11 |

| | | | | | | |
|---|---|---|---|---|---|---|
| | | BDV_P40 | pfam06407 | Borna disease virus P40 protein. This family consists of several Borna disease virus P40 | 83-306 | 9.33E-35 |
| SINV-16 | 1 | RNA_dep_RNAP | cd01699 | RNA_dep_RNAP: RNA-dependent RNA polymerase (RdRp) | 2679-2973 | 9.88E-54 |
| | | RdRP_1 | pfam00680 | RNA dependent RNA polymerase. | 2530-3014 | 1.94E-50 |
| | | rhv_like | cd00205 | Picornavirus capsid protein domain-like. Picornaviruses are non-enveloped plus-strand ssRNA | 610-804 | 2.12E-18 |
| | | RNA_helicase | pfam00910 | RNA helicase. This family includes RNA helicases thought to be involved in duplex unwinding | 1561-1664 | 2.05E-08 |
| | | rhv_like | cd00205 | Picornavirus capsid protein domain-like. Picornaviruses are non-enveloped plus-strand ssRNA | 341-530 | 2.68E-08 |
| | | CRPV_capsid | pfam08762 | CRPV capsid protein like. This is a family of capsid proteins found in positive stranded | 1094-1234 | 2.26E-04 |
| SINV-17 | 1 | RdRP_2 | pfam00978 | RNA dependent RNA polymerase; This family may represent an RNA dependent RNA polymerase. | 2182-2588 | 3.26E-88 |
| | | Viral_helicase1 | pfam01443 | Viral (Superfamily 1) RNA helicase. Helicase activity for this family has been demonstrated | 1619-1877 | 4.90E-28 |
| | | FtsJ | pfam01728 | FtsJ-like methyltransferase. This family consists of FtsJ from various bacterial and archaeal | 832-1012 | 7.14E-24 |
| | | Macro_Poa1p_like | cd02901 | Macro domain, Poa1p_like family. The macro domain is a high-affinity ADP-ribose binding module | 1935-2062 | 9.94E-18 |
| | | RlmE | COG0293 | 23S rRNA U2552 (ribose-2'-O)-methylase RlmE/FtsJ [ | 807-1014 | 4.93E-17 |
| | | Vmethyltransf | pfam01660 | Viral methyltransferase. This RNA methyltransferase domain is found in a wide range of ssRNA viruses | 90-418 | 1.11E-13 |
| | | rrmJ | TIGR00438 | cell division protein FtsJ; Methylates the 23S rRNA. | 832-1012 | 1.72E-09 |
| | | DEXQc_UvrD | cd17932 | DEXQD-box helicase domain of UvrD; UvrD is a highly conserved helicase involved in mismatch | 1610-1680 | 7.38E-05 |

| | | | | | |
|---|---|---|---|---|---|
| | recD_rel | TIGR01448 | helicase, putative, RecD/TraA family; This model describes a family similar to RecD | 1615-1726 | 5.31E-04 |
| | rrmJ | PRK11188 | 23S rRNA methyltransferase J; Provisional | 832-1006 | 6.77E-04 |
| | RecD | COG0507 | ATP-dependent exoDNAse (exonuclease V), alpha subunit, helicase superfamily I | 1615-1726 | 1.28E-03 |
| 2 | DiSB-ORF2_chro | pfam16506 | Putative virion glycoprotein of insect viruses. DiSB-ORF2_chro | 76-126 | 1.93E-13 |
| 3 | SP24 | pfam16504 | Putative virion membrane protein of plant and insect virus. SP24, | 66-210 | 2.06e-06 |

* No conserved domain was found according to CDD.

**Supplementary Table S4.** Data set used in the compositional analyses.

| Genus | Virus name | GenBank access number | | | | Host |
|---|---|---|---|---|---|---|
| | | RdRp | glycoprotein | nucleocapsid | NS4 | |
| *Phasivirus* | Badu virus | KT693187 | KT693188 | KT693189 | - | Insect |
| *Goukovirus* | Yichang Insect virus | KM817703 | KM817730 | KM817763 | - | Insect |
| | Gouleako virus | HQ541738 | HQ541737 | HQ541736 | - | Insect |
| *Phlebovirus* | Uukuniemi virus | D10759 | M17417 | M33551 | - | Vertebrate/tick |
| | Rift Valley fever virus | X56464 | M11157 | X53771 | - | Vertebrate/insects |
| | Munguba virus | HM566164 | HM566165 | HM566166 | - | Vertebrate/tick |
| | Toscana Virus | X68414 | X89628 | X53794 | - | Vertebrate/insect |
| *Orthobunyavirus* | La Crosse virus | U12396 | EF485031 | EF485030 | - | Vertebrate/insect |
| | Akabane virus | AB968525 | AB100604 | AB000851 | - | Vertebrate/insect |
| | Bunyamwera virus | X14383 | M11852 | D00353 | - | Vertebrate/insect |
| *Tenuivirus* | Rice hoja blanca tenuivirus | MG566074 | L54073 | L07940 | MG566077 | Plant/insect |
| | European wheat striate mosaic virus | MN044342 | MN044343 | MN044344 | MN044345 | Plant/insect |
| | Ramu stunt virus | KR094115 | - | - | - | Plant/insect |
| | Rice grassy stunt virus | AB009656 | AB010376 | AB000403 | AB010378 | Plant/insect |
| | Melon chlorotic spot virus | MH817469 | - | - | - | Plant/insect |
| | Rice stripe virus | D31879 | D13176 | X53563 | D10979 | Plant/insect |
| | Maize stripe tenuivirus | - | - | - | L13438 | Plant/insect |
| | Echinochloa hoja blanca virus | - | - | - | L48441 | Plant/insect |
| *Orthotospovirus* | Peanut bud necrosis virus | AF025538 | U42555 | U27809 | - | Plant/insect |
| | Tomato spotted wilt virus | D10066 | S48091 | D00645 | - | Plant/insect |
| | Groundnut ringspot virus | HQ644142 | MH742957 | MH742958 | - | Plant/insect |
| | Polygonum ringspot tospovirus | KJ541746 | KJ541745 | KJ541744 | - | Plant/insect |
| *Horwuvirus* | Wuhan horsefly Virus | KM817690 | - | - | - | Insect |
| Unclassified | S. invicta virus 14 | MT860236 - MT860243 | MT860244 - MT860251 | MT860252 - MT860259 | MT860260 - MT860267 | Insect |
| Unclassified | Fitzroy Crossing tenui-like | MT498812 | MT498813 | MT498815 | MT498814 | Insect |

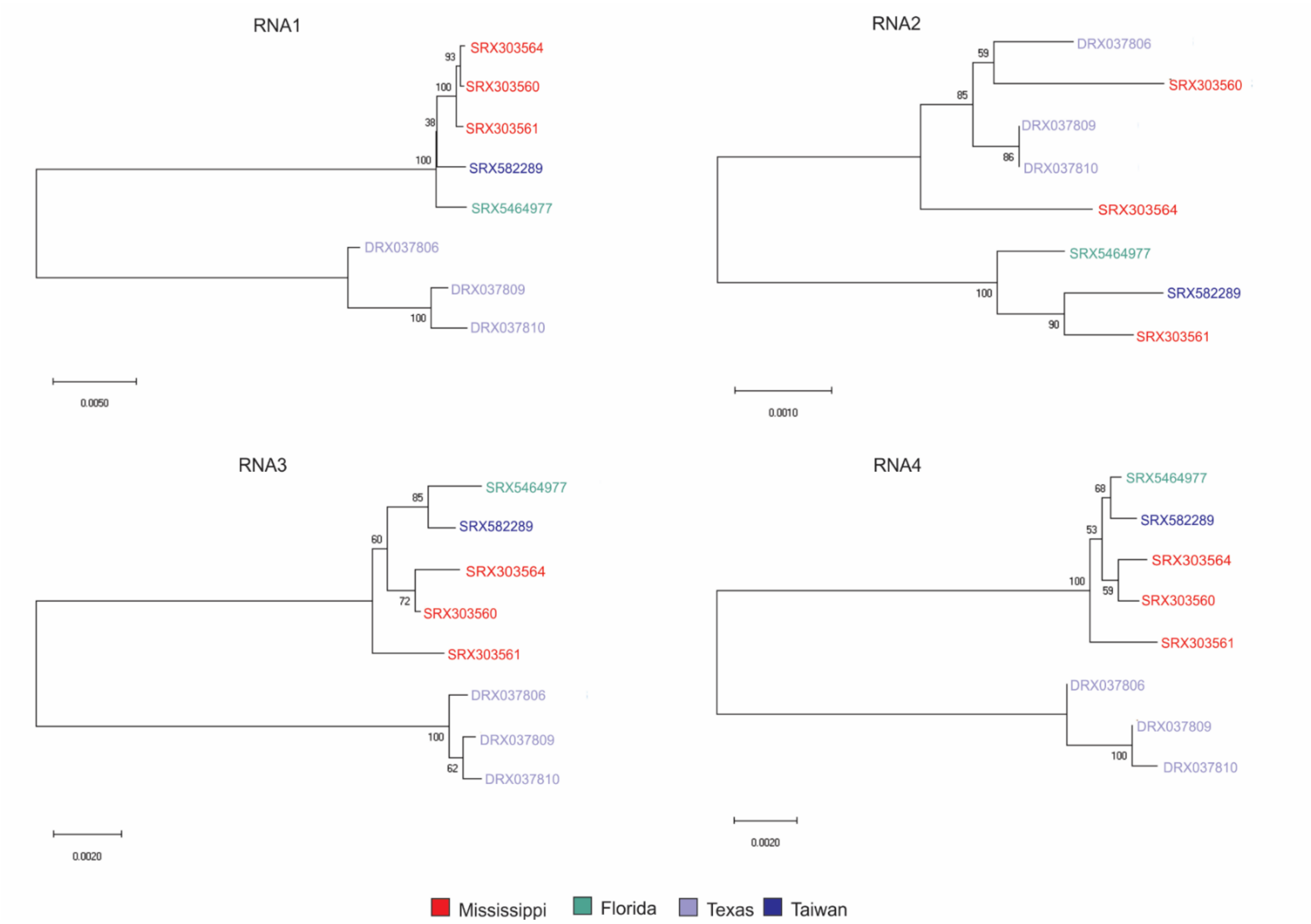


**Supplementary Figure S1.** Neighbor-Joining phylogenetic tree (1000 bootstrap replications) showing the clustering of the SINV-14 segments. Isolates are highlighted according to sampled place as indicated at the bottom of the figure.

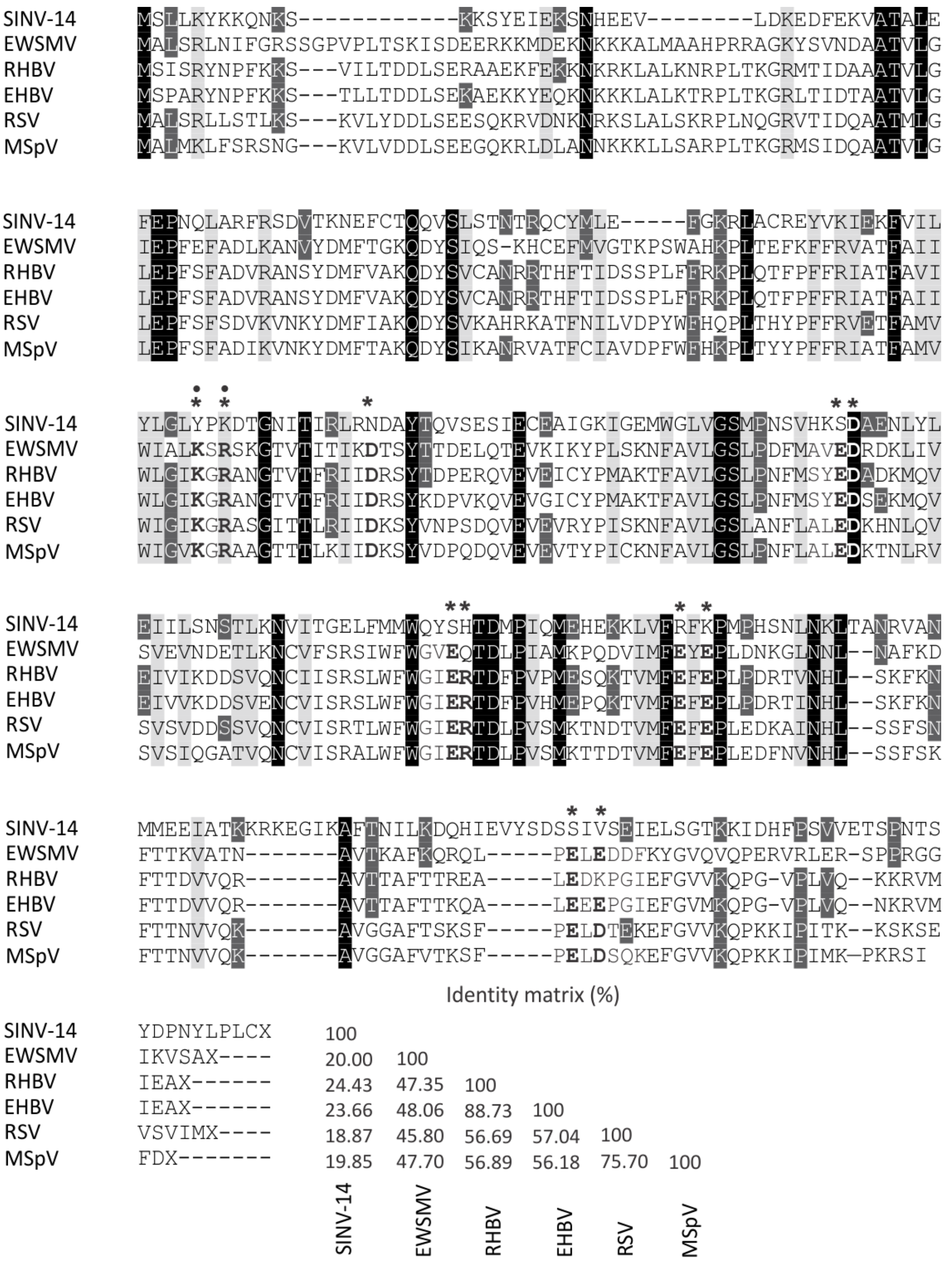


**Supplementary Figure S2.** Alignment of non-structural protein 4 (NS4) amino acid sequence of SINV-14 with those from typical plant tenuiviruses. Positions with single conserved amino acid are highlighted in black, groups of similar properties are highlighted in light gray and positions in

SINV-14 that shares the same amino acid with at least one plant tenuivirus sequence, are highlighted in dark gray. Sites containing residues essentials for cell-to-cell movement and long-distance movement are indicated by asterisk and filled circle, respectively. These sites were mapped using Maize stripe virus (MSpV) as a model (66). Matrix of pairwise identity of NS4 amino acid sequence are shown at the bottom of the figure. Multiple sequence alignment and pairwise identity were generated in Clustal O v. 1.2.4. European wheat striate mosaic virus (EWSMV), MN044345; Rice hoja blanca virus (RHBV), MG566077; Echinochloa hoja blanca virus (EHBV), L48441; Rice stripe virus (RSV), D10979; Maize stripe virus (MSpV), L13438.
